# ARCHER: Amortized cross-specimen pose estimation for cryo-electron microscopy

**DOI:** 10.64898/2026.08.21.746234

**Authors:** Nhan D. Nguyen, Bao Pham

## Abstract

Single-particle cryo-electronic microscopy (cryo-EM) pose estimation is traditionally solved anew for each dataset, where iterative refinement is done from scratch while the estimator learns to store the molecule in its weights. In this work, we show that pose inference is a generalizable, specimen-agnostic operation when conditioned explicitly on a reference volume. We introduce **ARCHER**, an amortized contrastive classifier that models the pose posterior over a discrete rotation grid. Trained across a variety of protein structures, it operates zero-shot *without retraining per structure*. This transferability is grounded in Fourier-space information mechanics, where all specimen dependence is captured by the reference structure’s power spectrum and spatial extent. ARCHER achieves a median angular error of 5.0^◦^ on 100 held-out test structures and 2.5^◦^ on experimental particles, matching dedicated estimators within 0.16 Å in 3D reconstruction. Crucially, downstream conformational signal is preserved. The leading conformational coordinate correlates at 0.97 with deposited benchmarks, faithfully reconstructing free-energy basins and mobile domains. These results overall demonstrate that cryo-EM pose estimation can be generalized across different structures.

## I. INTRODUCTION

Single-particle cryo-electron microscopy (cryo-EM) determines macromolecular structure from images of individual molecules frozen in vitreous ice^1,2^. In recent years, direct electron detectors and improved reconstruction algorithms brought the method to atomic resolution across a wide range of specimens^3–5^, and it is now the principal technique for assemblies that resist crystallization. However, the measurement is generally destructive and the electron dose is severely limited, so each particle is recorded at a signal-to-noise ratio well below unity and structure is recovered by averaging 10^5^ to 10^6^ of the raw particles.

Averaging requires knowing how each particle is oriented. Because the specimen is frozen in random orientations, the rotation that produced each image is a latent variable that must be inferred jointly with the structure^6–8^. By the central-slice theorem^9^, a projection samples the Fourier transform of the volume on a plane through the origin, where recovering the volume amounts to placing every measured plane correctly in three dimensions. Errors in that placement blur the average, and the placement problem dominates both the computational cost and the failure modes of the pipeline^10,11^.

### Iterative refinement

The established solution treats poses as missing data and marginalizes them. Maximum-likelihood formulations^6,12^ and their Bayesian extension^7^ alternate between computing a posterior over orientations for every particle and rebuilding the volume as a posterior-weighted backprojection, an expectation– maximization scheme^13^ implemented in RELION^10,14^, cryoSPARC^11^ and FREALIGN^15^. Reliability comes from evaluating half-sets independently and reporting a Fourier shell correlation between them^16,17^, with resolution read at a fixed criterion^18^ or one that accounts for the number of samples per shell^19^. This machinery is mature, and it restarts from scratch for every dataset: the scoring operation is recomputed from first principles each time, at a cost that scales with the product of particles and candidate orientations.

### Learned estimators

Neural approaches replace part of that loop. One family performs amortized inference within a single dataset of a single structure: cryoDRGN^20,21^ learns a conformational latent space with a coordinate-based decoder, while cryoAI and cryoFIRE^22,23^ add an encoder that predicts pose directly, removing the per-particle search. cryoSPIN^24^ refines an initial prediction by search, and other related works explore implicit^25,26^ and diffusion-based^27^ representations. A second family is supervised: given a solved structure and its assigned poses, a rotation classifier can be trained and applied to the same specimen, as in cryoPARES^28^ and earlier work on learned orientation assignment^29,30^. The CESPED benchmark^31^ standardized this setting. A third line targets ab-initio determination without any reference, either by amortized regression as in CryoFastAR^32^ or by classical common-lines synchronization^33–35^.

Across these families of approaches, the volume lives in the network parameters. An encoder trained on one specimen encodes that specimen’s projections, so a new protein requires new training, and the computation spent learning to compare a noisy image against a candidate view is discarded. Recent foundation-model work in cryo-EM has begun to target reusable components rather than per-dataset models: cryoFM^36^ learns a generative prior over densities, Cryo-IEF^37^ learns particle features by self-supervision, and learned regularization^38^ and map restoration^39,40^ transfer across specimens. Pose assignment itself has remained per-dataset. In this work, we instead ask whether the comparison at the heart of pose assignment can be learned once and reused.

The quantity that every pose estimator evaluates is the posterior *p*(**R** | *y, V*) over rotations, in which *y* ∈ ℝ^*L×L*^ is a single recorded particle image and *V* ∈ ℝ^*L×L×L*^ is the reference volume on a cubic grid of side *L* voxels. When *V* is supplied as an argument rather than fitted into the weights, this expression separates two quantities that per-dataset methods entangle. The first is the specimen, which is data and differs at every target. The second is the operation of matching a noisy projection to a candidate view, which is governed by the microscope – the contrast transfer function (CTF)^41^, the noise spectrum, and the slice geometry – and is therefore common to every specimen. Provided with the reference volume, the network we develop learns a matching function that transfers to molecules it has never seen, and such a reference volume is almost always available in practice, whether as a consensus map, a homologue, or a predicted structure^42–45^.

### ARCHER

Our approach, ***A***mortized ***R***eference***c***onditioned ***H*** ierarchical ***R***efinement realizes this posterior as a contrastive classifier over a discrete rotation grid, and we train it across 3,330 protein structures and apply it without retraining. It places particles at a median error of 5.0° on 100 structures held out of training, and at 2.5° on experimental data from EMPIAR-10076^46^. Its reconstructions come within 0.16 Å of per-dataset estimators where a target’s own sampling is the binding constraint. Conformational signal is left intact, in that the leading conformational coordinate recovered through our poses agrees with the one recovered through the deposited poses at a canonical correlation of 0.97.

We build a diffusion forward process that interpolates the identity into the CTF, so that a recorded particle is *exactly* its terminal state and denoising is one network evaluation at a known noise level rather than a sampled reverse trajectory – which is cheap enough to sit inside the training loop and differentiable through it. Its measured effect on pose accuracy is confined to fine precision on held-out synthetic proteins (see Sec. ( III D)). Every result reported here is from the three-channel configuration.

Lastly, we quantified the amortization capability of our network. Rotating a structure by a small angle displaces a Fourier component at radius *k* by an arc proportional to *k*, so the information about orientation grows as *k*^2^ times the spectral signal-to-noise ratio. This scaling follows from the microscope and from the geometry of the central-slice theorem, and the specimen enters it through its own power spectrum, which the reference volume supplies. That shared physics is what allows one encoder to learn from many maps, and it sets both the accuracy attainable and the point beyond which different methods converge on the same range of accuracy.

## II. RELATED WORK

### A. Orientation as a Latent Variable

The reconstruction problem was posed in its modern form by De Rosier and Klug^2^ and Crowther *et al*.^9^: each micrograph is a line integral through the specimen, so its Fourier transform is a central section of the transform of the volume, and the structure follows once enough sections are placed correctly. With orientations unknown, placement and reconstruction are coupled.

Two classical strategies resolve the coupling. Projection matching scores every particle against reference projections of a working volume and iterates^12,15^, performing an exhaustive search with a whitened matched filter and an explicit false-positive threshold, which detects and orients individual molecules in crowded images^47,48^. This is the classical pipeline we compare against throughout.

Meanwhile, maximum-likelihood formulations instead treat the pose as a latent variable and marginalize it^6^, which Scheres^7^ casts in a Bayesian framework with a regularizing prior on the volume and implemented as expectation-maximization^13^ in RELION^10,14^; cryoSPARC added stochastic gradient descent for initialization and branch-and-bound search^11^, later with adaptive regularization^49^. A third route avoids a reference entirely by exploiting the common-line geometry of pairs of projections and solving the resulting synchronization problem^33–35^.

Overall, these methods share a computational signature. The pose posterior is recomputed from first principles for every particle of every dataset, at a cost proportional to the number of candidate orientations, and nothing learned on one specimen is transferrable to another specimen.

### B. Conformational Heterogeneity

Most interesting specimens are flexible, and the pose problem is entangled with a conformational one. Discrete treatments assign particles to a small number of classes^50,51^, while multi-body refinement partitions the molecule into rigid units with independent orientations^52^. Continuous treatments learn a low-dimensional latent space: cryoDRGN couples an encoder to a coordinate-based volume decoder^20,21^, 3DFlex models deformation fields directly^53^, and RECOVAR^54^ estimates the covariance of the volume distribution and deconvolves the per-particle posterior to recover a conformational density. The last property matters for evaluation: because RECOVAR returns an embedding with per-particle uncertainty and is deterministic given its inputs, running it twice on identical particles under different pose estimates isolates the effect of the poses, which is the comparison we use in Sec. (IV C).

Heterogeneity also sets the difficulty of pose assignment itself. A flexible molecule presents projections that no single rigid volume explains, so scoring against a consensus map is systematically mismatched – the regime in which estimators differ most.

### C. Learned Pose Estimation

#### Per-structure Encoders

Following the amortized-inference pattern of variational autoencoders (VAEs)^55^, cryoAI and cryoFIRE replace the per-particle search with a VAE trained jointly with a volume representation^22,23^, and cryoSPIN adds a search-based correction to the encoder’s prediction^24^. These methods amortize over the particles of a single dataset pertaining to a structure, so the training cost is repaid across images of the same specimen. Replacing search with a forward pass trades exactness for speed, and the resulting amortization gap between the inferred posterior and the true one is a known cost of the design^56^. Because the volume lives in the decoder, the encoder is meaningful only for the targeted dataset it was fit on.

#### Supervised, Structure-specific

Given a solved structure and its assigned poses, orientation assignment becomes supervised learning on SO(3)^29,30^, standardized by the CESPED benchmark^31^ and carried to production accuracy by cryoPARES^28^. This family learns the same matching operator we do, but ties it to one molecule, so its cost recurs per specimen: training it on four CESPED targets took 90.5 single-GPU hours here, against a single training run that serves all of them (see Tab. S14 in the Appendix).

A protocol difference is worth naming before any of these are compared. The cryoDRGN workflow, and its ab-initio successor^57^, interpose an interactive curation step in which particles judged to be junk are filtered out before or between reconstruction rounds^20,21^. In contrast, our approach is an end-to-end process: every particle is scored, none is removed, and no manual intervention enters between the particles and the poses. Comparisons that hold the particle set fixed, as ours do, are therefore stricter on us than on a curated pipeline.

#### Reference-free Amortization

CryoFastAR regresses poses directly from particle sets without a reference^32^, and related work explores implicit representations^26^ and diffusion models^27,58^ for ab-initio determination. cryoDRGN-AI is the current state of the art in this setting, recovering both structure and motion without a starting model^57^, and CryoBench supplies the standardized heterogeneity datasets on which such methods are now compared^59^. This is the hardest setting and the one where heterogeneity costs most (see our comparisons in Sec. IV).

#### Representing distributions on SO(3)

All learned estimators must place a distribution on a curved space. Equivariant architectures^60,61^ and the Image2Sphere^62^ construction build the symmetry into the network, while implicit-PDF^25^ represents the density by evaluating an energy at sampled rotations. We take the discretization route, with a HEALPix grid^63,64^ that gives near uniform coverage and a controllable spacing, and recover sub-cell precision by a continuous refinement step rather than by a finer grid.

### D. Reusable components

Parts of the pipeline have already been shown to transfer across specimens. Particle picking^65^ and micrograph preprocessing^66^ are routinely handled by models trained once and applied broadly. On the map side, learned regularization inside refinement^38^ and post-hoc restoration^39,40^ generalize to structures never seen in training, and cryoFM learns a generative prior over densities usable as a plug-in for downstream tasks^36^. That prior is defined over maps rather than over orientations, so cryoFM contributes to a pipeline at the reconstruction and restoration stages and produces no per-particle pose; it therefore appears here as a map restorer (Sec. IV B) and cannot be placed on the pose accuracy axis at all, which is a difference in what the two models represent rather than a comparison either way. Cryo-IEF learns particle-level features by self-supervision across many datasets^37^, demonstrating that image statistics transfer even when structures do not.

Orientation assignment has stayed outside this trend, and the reason is structural rather than incidental: the quantity a pose estimator must know is the specimen, and every design so far has supplied it by fitting. Providing it as an argument is what makes the remaining operation – the comparison of a noisy projection against a candidate view under a known transfer function – shared across specimens, and therefore learnable once. Secs. (III) and (IV) make that statement quantitative and show what it implies for the accuracy attainable.

## III. METHODS

### A. Forward model

Volumes are sampled on a cubic grid of side *L* voxels and images on the corresponding *L v L* grid, following the discretization used in the mathematical cryo-EM literature^8,67^; *L* is reserved for this box side throughout and *C* for the number of input channels of a network. Table S1 in the Supplementary Material displays every symbol with the corresponding space.

Let *V* ∈ ℝ^*L×L×L*^ be a volume, 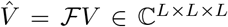 its discrete Fourier transform, **R** ∈ SO(3) a rotation, and *ξ* ∈ ℝ^3^ the per-particle contrast-transfer parameters (defocus along the two astigmatic axes and the astigmatism azimuth; voltage, spherical aberration and amplitude contrast are fixed per dataset). A particle image *y* ∈ ℝ^*L×L*^ is modeled as

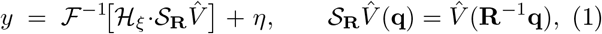

where *S*_**R**_ : ℂ^*L×L×L*^ → ℂ^*L×L*^ is the central-slice oper-ator, _*ξ*_ ∈ R^*L×L*^ the contrast transfer function (CTF), **q** ∈ ℝ^3^ a Fourier coordinate restricted to the slice plane 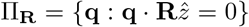, and *η* ∈ ℝ^*L×L*^ colored Gaussian noise whose power in the shell at radius *k* = ∥**q**∥ is *σ*^2^(*k*), measured from real micrographs. The transfer function is astigmatic, with the sign convention of Ref.^41^. Equation (1) is the flat Ewald-sphere approximation^68,69^; it is exact to the resolutions considered here and it makes handedness an exact degeneracy, since *V* and its mirror produce identical projection sets^70^.

Because the noise is independent between Fourier coefficients, the log-likelihood of a rotation given one particle is a weighted least-squares residual,

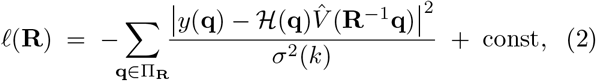

and every estimator compared in this work is, explicitly or implicitly, an attempt to maximize or to marginalize Eq. (2).

### B. Information geometry of orientation estimation

Two questions decide whether one estimator can serve many specimens: how much orientation information a single particle carries, and how much of that information depends on which molecule is in the ice. Both follow from Eq. (2).

#### a. The derivative of a slice under rotation

Perturbations of **R** live in the Lie algebra so(3). Take the one-parameter family **R**(*δ*) = **R** exp(*δ ê*^∧^) for a unit axis *ê* ∈ S^2^, an angle *δ* ∈ ℝ and the skew-symmetric generator *ê*^∧^ ∈ so(3). Writing **p** = **R**^−1^**q** and differentiating the sampled coefficient at *δ* = 0,

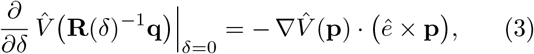

whose second factor is the velocity of the sampling point under the rotation, of magnitude

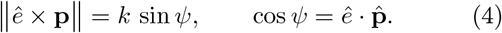

A rotation by *δ* transports a component at radius *k* along an arc of length *kδ* sin *ψ*, so the same angular perturbation moves high-frequency content proportionally further. This displacement is the geometric origin of everything that follows, and the classical relation between angular accuracy and attainable resolution rests on it^18^.

#### b. Orientation information grows as k^2^ SSNR(k)

For complex Gaussian noise the Fisher information of *δ* is the inverse-variance-weighted squared derivative of the model,

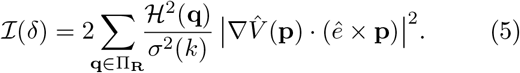

Substituting Eq. (4) and grouping the sum into shells gives the contribution of shell *k*,

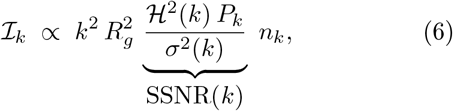

in which *P*_*k*_ is the signal power in shell *k, n*_*k*_ ∈ ℕ the number of Fourier samples it holds, SSNR(*k*) the spectral signal-to-noise ratio (SSNR) of that shell, and *R*_*g*_ the radius of gyration of the density about the axis of rotation. Three reductions carry Eq. (5) to Eq. (6), each an approximation rather than an identity. 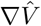 is the transform of −2*πi* **x***V* (**x**), so its shell power is of order 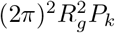 rather than *P*_*k*_; this moment approximation is what carries the specimen’s spatial extent into the result. The gradient is then taken isotropic within a shell, which replaces the projection onto one direction by 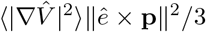 and leaves a constant from averaging sin^2^ *ψ*. And *V* is real, so summing over the full slice counts each independent coefficient twice, a factor of about two absorbed into *n*_*k*_. The factor 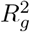 is not a nuisance constant: a larger particle carries its high-frequency content through a longer arc for the same rotation, so orientation is easier to determine. Read in real space, a point at radius *r* moves by *rδ*, and requiring that to stay within a resolution element *d* gives *δ* ≲ *d/r* — the same inverse relation between angular accuracy and attainable resolution that Rosenthal and Henderson write as *δ* ≲ *d/D* over a particle of diameter *D*^18^.

Two multiplicities compound in Eq. (6): the arc length grows as *k*, contributing *k*^2^ after squaring, and the shell population grows as *n*_*k*_ ∝ *k*^2^. Orientation evidence is therefore concentrated at high spatial frequency, where the SSNR is smallest. The useful band is the maximizer of *k*^2^ SSNR(*k*), and that competition — geometry pushing up, radiation damage and the envelope pushing down sets a finite optimal resolution range for pose assignment.

#### c. The prediction, and what the data give

Scoring each shell by a separately normalized correlation removes *n*_*k*_ and the shell’s absolute power, leaving a curvature Λ_*k*_ whose remaining specimen dependence runs through the correlation *c*_*k*_ that shell attains at the true pose. It does not remove 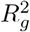, a property of the molecule rather than of the shell; that factor is constant here because the exponent is fitted within a single specimen, where it enters the intercept and not the slope. What remains is a prediction with no free parameter,

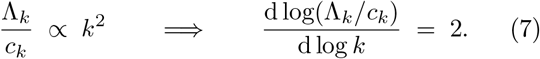

Measured on real particles at their deposited orientations — the Fisher information is defined at the true parameter — the curvature follows Λ_*k*_*/c*_*k*_ ∝ *k* ^1.89^ over shells *k* = 4 to 32, spanning 0.90 decades (Fig. 9A). Neighboring shells are correlated, and the fit residuals carry a lagone autocorrelation of 0.40, so the ordinary least-squares standard error of 0.037 is optimistic; the autocorrelation-robust value is 0.042, placing the measurement 2.6*σ* below 2. That deficit is not a bend in the power law, since adding a quadratic term in log *k* improves the fit by 0.5*σ*. Two effects depress the slope uniformly and survive the *c*_*k*_ normalization: a residual envelope or *B*-factor, and noise bias in *c*_*k*_ at the outer shells. We read the measurement as agreement to within a few percent rather than exact confirmation. Either way the scaling is obeyed by the data with no network involved.

#### d Bound on attainable accuracy

Summing Eq. (6) over shells and inverting gives the Cramér–Rao bound^71,72^ for an unbiased estimator of a single rotation component,

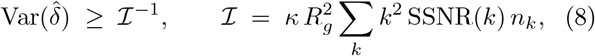

with *κ* a purely geometric factor collecting the shell averages and the Hermitian double-count above. Three consequences follow. Accuracy improves through shells that carry signal, so band-limiting a particle below the peak of *k*^2^SSNR discards orientation information irreversibly. The bound is per particle and independent of the estimator, so a gap between Eq. (8) and a measured error is a property of the algorithm. And because SSNR is fixed by the sampling of a given dataset, Eq. (8) is the formal statement of the specimen-level ceiling observed in Sec. ( IV D).

Two qualifications set its scope. Equation (8) bounds one rotation component with the other two known, and is optimistic by a factor of order unity against the corresponding element of the inverted 3 × 3 information matrix; *κ* also varies with the axis, since a rotation about the beam keeps the sampling point in the slice plane at sin^2^ 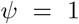 while a perpendicular axis averages to 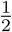, a factor 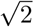 in angular standard deviation that the three body-axis probes of Eq. (17) see in aggregate. The bound also governs the local basin around the true pose, and says nothing about assigning a particle to the wrong basin. That second failure is a detection question, set by whether the matched-filter statistic *ρ*^2^ clears the 2 ln|*G*| threshold imposed by the competing candidates — the multiple-hypothesis accounting that 2D template matching applies to a whitened matched filter searching orientations exhaustively against a reference^47,48^. Neighboring grid poses give correlated scores, so the effective number of independent candidates is smaller than |*G*| and that threshold is conservative here. A local precision bound and a global detection criterion together determine attainable accuracy.

#### e. Why one estimator serves many specimens

Write the inference we wish to perform as a map from data and reference to a distribution on rotations,

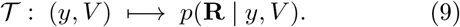

Bayes’ rule with Eq. (2) gives *p*(**R** *y*,| *V*) ∝ exp *ℓ*(**R**; *y, V*), so *T* is determined by three objects: the transfer function, the noise spectrum, and the slice geometry of the central-slice theorem. None of them depends on which molecule is in the ice. The specimen enters Eq. (6) through *P*_*k*_ and its spatial extent *R*_*g*_, both of which the reference supplies at inference time.

A per-dataset estimator learns instead the partially applied map *T*_*V*_ : *y* → *p*(**R** | *y, V*) with *V* absorbed into the parameters. The two differ in what generalizes. Fitting *T*_*V*_ for *M* specimens requires *M* independent fits, each using only that specimen’s particles; fitting *T* once uses all particles from all specimens for the single operator that is common to them, and treats *V* as a nuisance argument redrawn at every training step: each step supplies one reference volume together with a batch of *B* particles generated from it, so a single template bank is rendered once and shared by the whole batch (Sec. III F). Training across many maps is then a variance-reduction device for the shared operator rather than a compromise between specimens.

#### f. Discretization floor

The posterior is evaluated on a finite grid *G* ⊂ SO(3), so even an exact scorer inherits a quantization error. For |*G*| cells covering SO(3), whose Haar volume is 8*π*^2^ in the metric where geodesic distance is the rotation angle, the mean nearest-neighbor spacing scales as

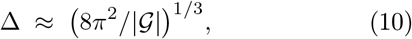

giving 7.4° at the |*G*| = 36,864 used here, against a measured median nearest-neighbor spacing of 7.40°. Reaching below this floor requires a continuous step, which motivates the refinement of Sec. (III E); conversely, refining a grid whose spacing is already below the Cramér–Rao scale of Eq. (8) buys nothing.

### C. Reference-conditioned posterior

Templates 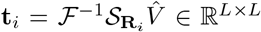 are rendered from the reference by central-slice extraction on a HEALPix grid *G* ⊂ SO(3)^63,64^ and encoded once per training step; particles are encoded per image. Both encoders are residual convolutional networks^73^ mapping into the unit sphere S^511^ ⊂ ℝ^512^ (10.4 M parameters per branch): the particle encoder *f*_*θ*_ : ℝ^*C×L×L*^ → S^511^ takes the *C*-channel image stack and the template encoder *f*_*ϕ*_ : ℝ^*L×L*^ → S^511^ a single rendered slice. Pose logits are

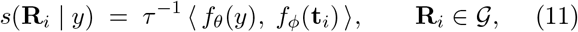

with *τ >* 0 a learned temperature^74,75^ that falls from 0.07 at initialization to 0.0212, sharpening the posterior by a factor of 3.3. The objective is cross-entropy against a geodesic soft target 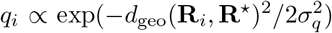 with *σ*_*q*_ = 6°, which supplies gradient to neighboring cells and respects the metric of SO(3). Encoder topology, the hard-negative pool and the optimizer settings are given in App. B.

The CTF enters as image content: the *C* channels carry the raw particle and its phase-flipped counterpart, and templates are rendered without a transfer function. This costs information and buys amortization, and both sides are quantifiable. The sufficient statistic of Eq. (2) is ⟨ℋ · **t**, *y*⟩ */σ*^2^, which weights each shell by the transfer function’s amplitude; phase flipping keeps the sign and discards that weighting. Writing ⟨·⟩_*w*_ for the average under the weight *w*_*k*_ = *P*_*k*_*/σ*^2^(*k*), the ratio of the two informations is

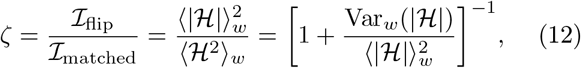

so the loss is the relative variance of |ℋ| across the band, and *ζ* ≤ 1 by Cauchy–Schwarz with equality for a flat transfer function.

Evaluated over the real per-particle defocus values of EMPIAR-10409 at the measured noise spectrum, *ζ* = 0.542 ± 0.009: the phase-flipped scorer retains 54% of the available orientation information, a factor 1.36 in angular standard deviation. What it buys is that templates are encoded once per specimen rather than once per particle, since applying *H* inside the similarity would make the template embedding defocus-dependent. That is the difference between one template bank per specimen and one per particle, and it is what makes amortization affordable.

### D. Diffusion denoising channel

At the signal-to-noise ratios of a single particle, the shells that carry orientation information by Eq. (6) are the ones the noise dominates, so restoring amplitude there is the operation that stands to sharpen a pose. The diffusion channel performs it: a Fourier-domain denoiser, trained jointly with the pose objective, supplies a denoised copy DN(*y*) of the particle as a third input channel to the encoder. The construction turns on one observation. In a diffusion model the data sit at *t* = 0 and noise is added to reach the terminal state; here we choose a forward process whose terminal state *is* the measurement, by interpolating the identity into the transfer function,

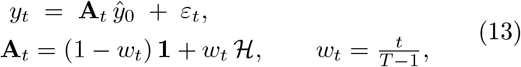

where *t* ∈ {0, …, *T* − 1} indexes the noise level, *T* ∈ ℕ is the number of levels, *ŷ*_0_ ∈ ℂ^*L×L*^ is the clean Fourier-domain image, **A**_*t*_ ∈ ℝ^*L×L*^ is the transfer operator acting elementwise, **1** ∈ ℝ^*L×L*^ is the all-ones array, *w*_*t*_ ∈ [0, 1] is the interpolation weight, and *ε*_*t*_ ∈ ℂ^*L×L*^ is zero-mean Gaussian of variance 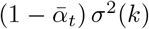, with 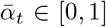 following a cosine schedule^76,77^. Because a recorded particle is formed as ℋ *ŷ*_0_ + *η*, Eq. (13) places it at *t* = *T* − 1 *exactly* : the denoiser is evaluated once at a known level rather than sampled over a reverse trajectory, which is

what makes it cheap enough to sit inside the training loop and differentiable through it. Equation (13) carries no 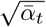 factor on the signal and its terminal marginal is the measurement rather than a fixed Gaussian, which places it in the corruption-schedule family, where the forward process is a physical degradation and the terminal state is the observation^78–82^.

A U-Net *ε*_ϑ_^83^, with parameters *ϑ* distinct from those of the two encoders, predicts the noise from the real and imaginary parts of *y*_*t*_, the transfer function and a sinusoidal time embedding. It is trained with the *ε*-prediction objective^76^, weighted by the inverse noise variance so that every shell contributes on the scale of its own uncertainty,

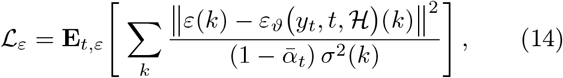

and the total objective is ℒ = ℒ_CE_ + ℒ_*ε*_, so two gradients reach this network: the *ε*-prediction loss with *t* drawn uniformly, and the pose cross-entropy through the pipeline. Given 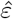, the clean estimate follows by Wiener inversion^84^,

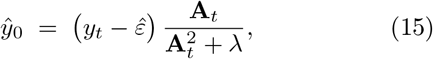

with 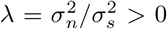 the ratio of leftover-noise to signal power. The gain is the linear estimator 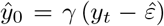 minimizing **E** |*ŷ*_0_ − *y*_0_|^2^ when the residual uncertainty is treated as a signal of prior power 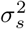 against leftover noise of power 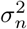, which gives 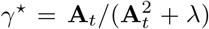. The regularizer is what makes Eq. (15) usable: direct inversion by **A**_*t*_ diverges at the zeros of *ℋ*, whereas this gain is bounded by 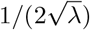 at 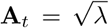 and falls to zero as **A**_*t*_ → 0^84^, so the estimator declines to invent content in the transfer function’s blind bands.

The channel acts on fine precision. Over 150 step-matched held-out evaluations on synthetic proteins, the median angular error is 5.07° against 5.22° and the fraction within 5° is 0.493 against 0.474, leading at 122 and 121 of the 150 evaluations, while the fraction within 15° is unchanged at 0.755 against 0.753: the denoised channel sharpens the estimate inside the correct basin and preserves the coarse assignment. On experimental particles, we reached pose median on EMPIAR-10409 to 0.01° and the reconstructions to 0.001 Å. The results reported throughout this work are from the three-channel configuration, CTF-modulated particles, phase-flipped, and the denoising channel.

### E. Hierarchical refinement

Inference proceeds from the grid maximum to a continuous estimate. Given current poses, a volume is rebuilt by Wiener-filtered backprojection, every particle is re-scored against re-rendered templates by a shell-normalized correlation restricted to low shells, and each pose is polished by a derivative-free Newton step.

Let *s*(**R**) be the score of Eq. (11) or its classical counterpart. Parameterizing a neighborhood of the current estimate by the exponential map, 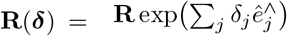 with ***δ*** ∈ ℝ^3^, a second-order expansion gives

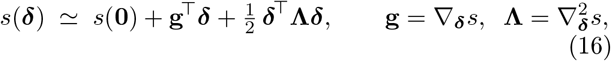

with gradient **g** ∈ ℝ^3^ and Hessian **Λ** ℝ^3*×*3^, whose stationary point is ***δ***^⋆^ = −**Λ**^−1^**g**. The update is applied on the group, **R** ← **R** exp(***δ***^⋆∧^), so the estimate never leaves SO(3).

We evaluate **g** and the diagonal of **Λ** by central differences, which needs no gradient through the scorer, so the same routine refines the learned posterior and the classical matched filter. Writing 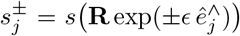 and *s*^0^ = *s*(**R**) for the score probed along the three body axes *ê*_*j*_ ∈ S^2^, *j* = 1, 2, 3, at probe angle *ϵ >* 0,

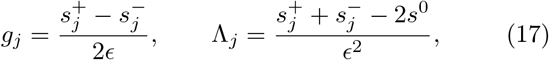

are the directional first and second derivatives, and the update **R** ← **R** exp(***δ***^∧^) with *δ*_*j*_ = *g*_*j*_*/* Λ_*j*_ is taken where Λ_*j*_ *<* 0 and clamped to 1° per axis otherwise. Where the curvature has the wrong sign the step reverts to that clamped gradient move, which keeps the update stable on the flat plateaus between grid cells. The probe scale is set by the two angular scales the problem supplies: it must exceed the noise-induced roughness of the score and stay below the curvature scale of its peak, and *ϵ* = 1° against a grid spacing of 7.4° satisfies both (Fig. 9C). The diagonal of the Hessian is evaluated at six renders per step; the full 3 × 3 would cost six more.

Two loops are nested here and are counted separately in Alg. 2. The *outer* loop rebuilds the volume, re-scores

#### ALG. 1.

Training. One step is one specimen; the reference is data rather than a parameter, so a single set of weights is fit across the bank.

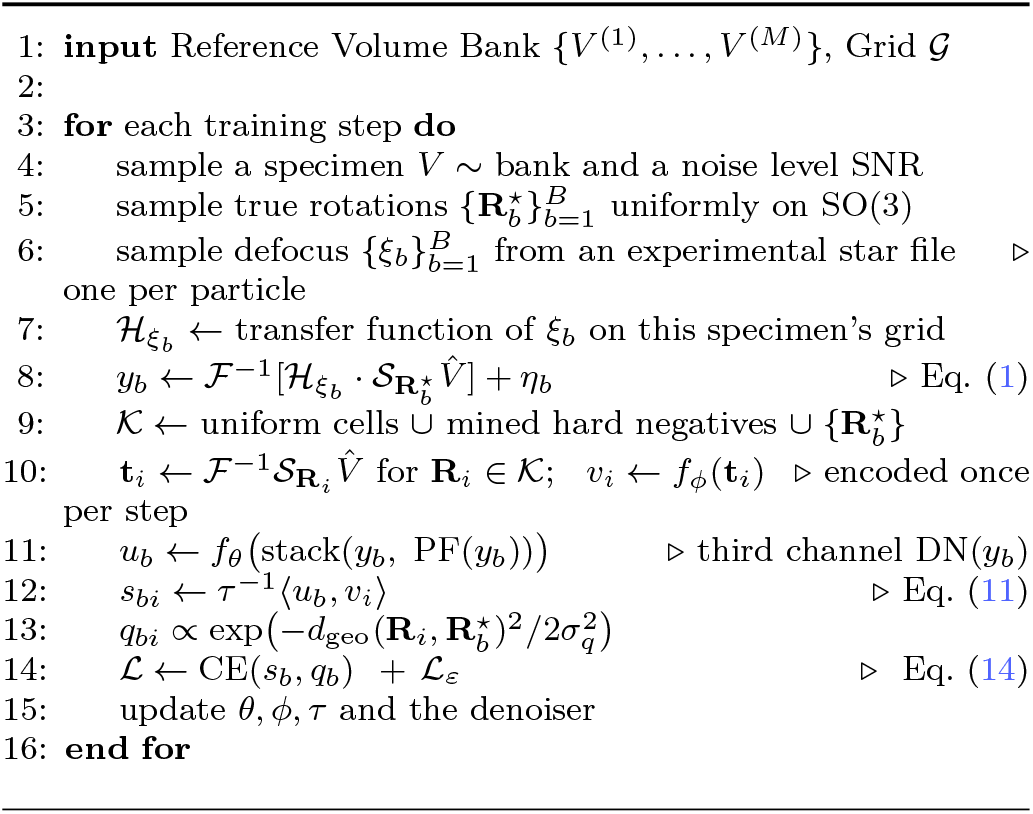

against it and refines, and runs *T*_iter_ = 3 times; the *inner* Newton loop takes *T*_newton_ = 4 update steps within each of those, at fixed volume and fixed templates. Half-sets are processed independently throughout — each half is backprojected into its own volume and every particle is scored only against the volume built from the opposite half — preserving the gold-standard separation^16,17^.

### F. Data

#### Volumes and Particles

Training particles are sampled from the training datasets (pertaining to different structures) using the known pose information relative to each structure. Our training sets are from the Electron Microscopy Data Bank (EMDB)^85^, where we used 3,330 structures while 100 structures are held out for evaluation. For each volume of each structure, *training episodes* are drawn from those 3,330 structures during training. Here, a training episode is a volume and 96 particles generated through the forward model of Sec. (III A): a central slice of the volume, a CTF whose defocus is drawn

##### ALG. 2.

Hierarchical inference. Half-sets are processed independently, so no particle contributes to the reference against which it is scored.

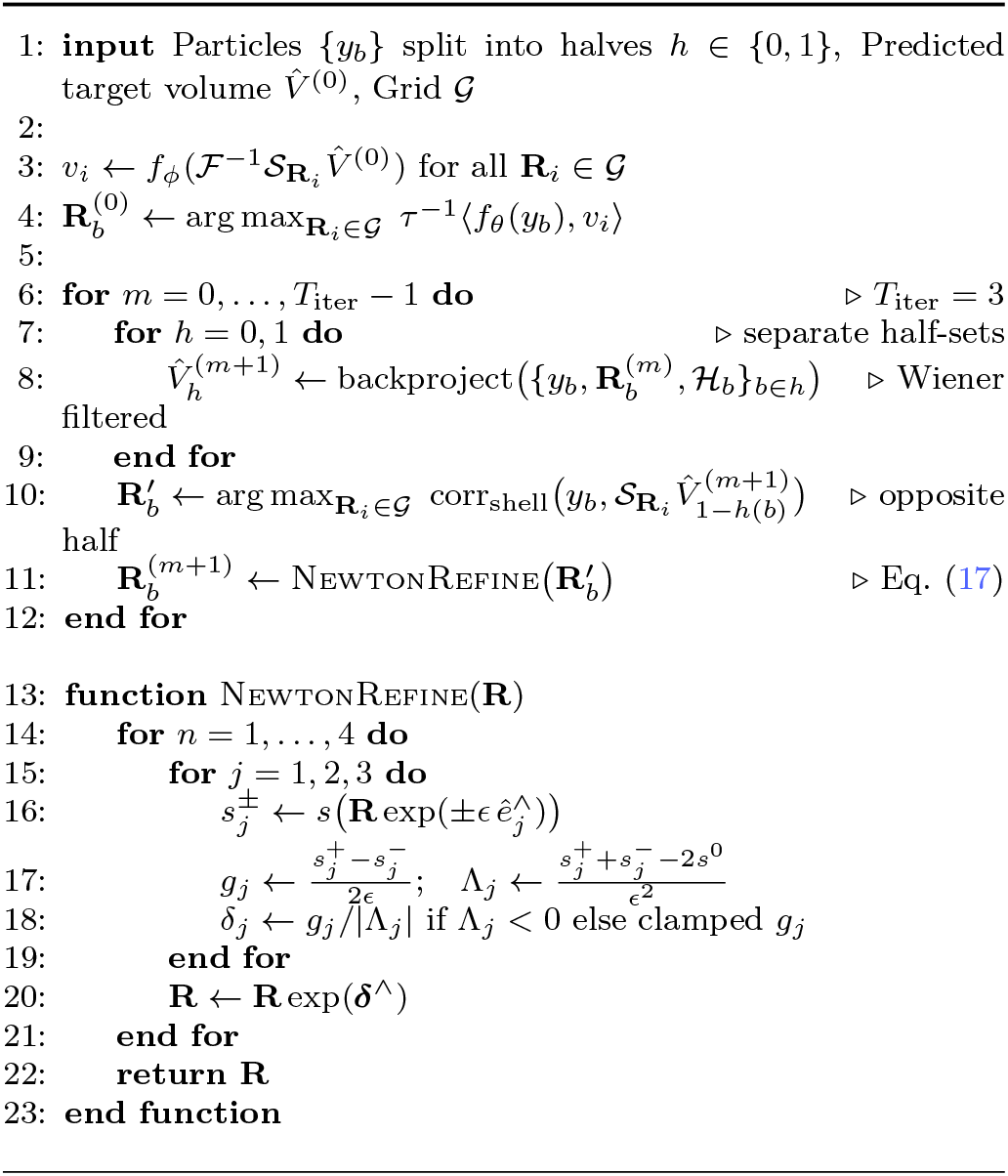

from the corresponding training set, and additive noise colored by a power spectral density measured from real micrographs rather than white, at signal-to-noise ratios sampled from 0.005 to 0.075. Training runs for 2 × 10^6^ of these episodes.

Moreover, for the volumes, we used the deposited half-maps, whose native boxes span 94^3^ to 268^3^ at roughly 1.5 Å per voxel. Each half-map is reduced to a common 64^3^ box by cropping in Fourier space rather than in real space: a real-space crop would truncate the larger proteins, whereas discarding high frequencies retains the whole molecule at coarser sampling, which is the operation the resolution argument of Sec. (III B) assumes. Because the crop is to a fixed box rather than a fixed sampling, voxel size varies with the structure (with median value of 4.7 Å alongside 90% of structures between 3.1 and 6.9 Å), so the model meets a range of scales during training.

### G. Evaluation

The held out structures are scored on simulated particles, which isolates generalization to unseen proteins from the change of domain. Experimental performance is measured, without retraining, on deposited data from the Electron Microscopy Public Image Archive^46^: EMPIAR-10076 for pose accuracy and heterogeneity, and the CESPED benchmark targets^31^ EMPIAR-10166, 10280, 10409, 10648 and 11120 for reconstruction. Training volumes come from the EMDB and evaluation particles from EMPIAR, but the deposited structures of the benchmark targets are not among the volumes trained on. Meanwhile, the inference pipeline, detailed in Alg. (2), takes in the predicted volume of the target structure 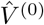, and iteratively refines the pose information (to refine the volume) for *T*_iter_ iterations.

Angular error is the geodesic distance on SO(3) to the deposited pose, reported as a median and as the fraction within a threshold. Map metrics follow the CESPED protocol^31^: half-sets are reconstructed independently with RELION^14^ at the predicted orientations and compared to the deposited structure inside the benchmark mask, with resolution read at the 0.143 criterion^18^. Conformational agreement is the canonical correlation between RECOVAR embeddings^54^ of identical particles differing only in rotations, with a shuffled-pose floor measured through the identical pipeline. Restoration is evaluated with EMReady^39^ and cryoFM^36^ using their published metrics. See Figs. (5) and (6) for the benchmark results.

Conformational landscapes are compared through those same embeddings. RECOVAR’s deconvolved density *p*(*z*)^54^ reads as a free energy through −log *p*(*z*), and the local maxima of *p* are the *basins*, the stable states. Comparing the map at one basin with all the others would treat a difference across a high barrier the same as one the particle crosses freely, so each destination is weighted by how likely the particle is to reach it,

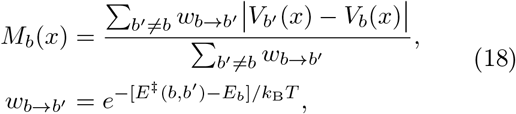

with *V*_*b*_ the map at basin *b* and *E*^*‡*^ the lowest high point on any path joining two basins — the minimax barrier used to organize energy landscapes into disconnectivity trees^86^, with the exponential weight the Arrhenius factor for escape over it^87^. This *mobility* is large where a structure differs from the states the particle can actually reach. Two pose sets are compared by matching their basins on map correlation and then comparing the mobility fields at the matched basins.

## IV. RESULTS

### A. Pose information prediction

ARCHER embeds a particle and a bank of reference templates into a shared unit sphere and reads the pose off the maximum of a temperature-scaled cosine similarity (see Fig. 2 and Alg. 1). The rotation grid, for the reference volume, is a HEALPix discretization of SO(3)^63,64^ with 36,864 cells at a median spacing of 7.4°.

**FIG. 1.**
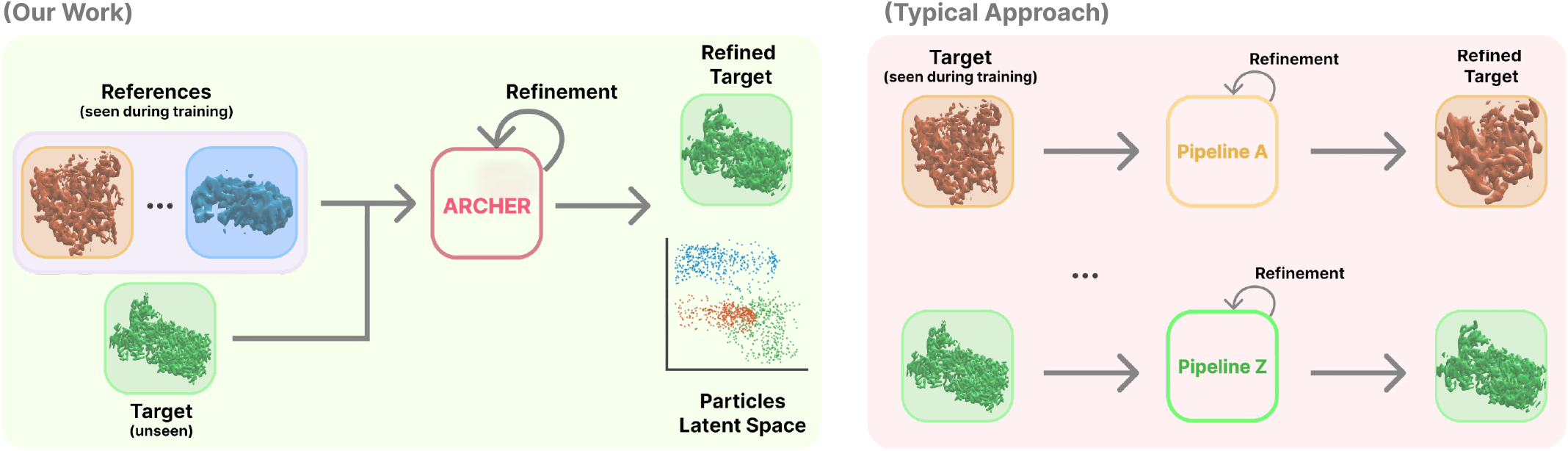
Comparison between ARCHER and typical approaches. ARCHER takes the reference volume as an *argument*. It is trained once across many structures and then applied to a target it has never seen, whose reference is handed to it at inference time alongside the particles. Refinement is applied afterwards on the estimate it produced, where the particle embeddings it computed support downstream heterogeneity analysis. In other methods, the specimen is absorbed into the weights, so each target requires its own fitted model and its own training run, and nothing learned on one target is transferred to the next.

**FIG. 2.**
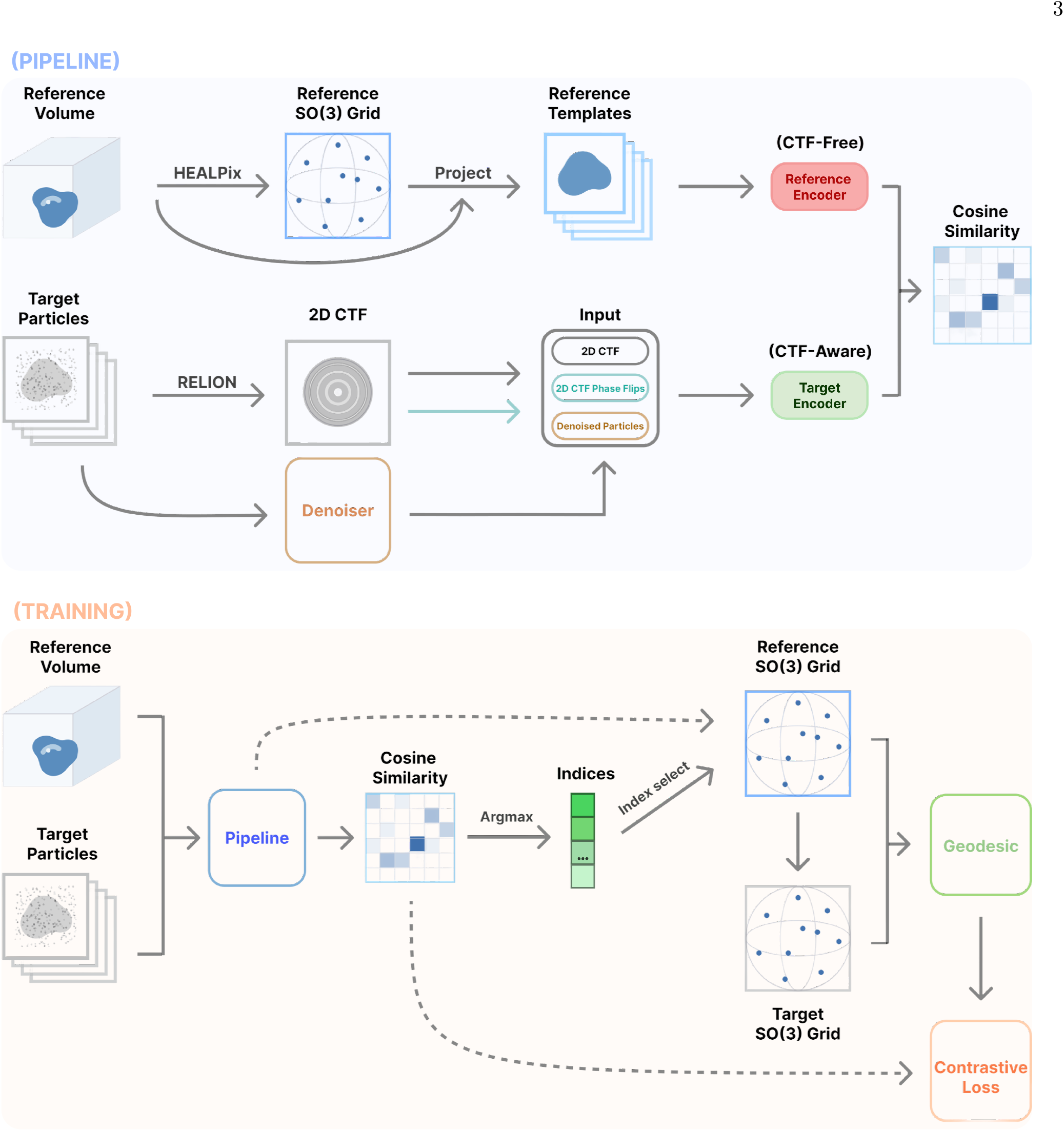
Overview of ARCHER’s architecture. *Pipeline*: The reference volume is sliced at every cell of a HEALPix grid on SO(3), giving a fixed bank of templates that is embedded *once* per specimen. Each particle is embedded by a second encoder from a stack of image channels: the recorded particle, its phase-flipped copy, and denoised particles. The transfer function enters the network through the pixels it modulates. The pose **R** is the grid cell whose template embedding maximizes a temperature-scaled cosine similarity against the particle embedding. *Training* : The same forward pass is run on simulated particles whose true rotation is known. That rotation is mapped to its nearest grid cell, and the resulting cosine similarities are trained against a target smoothed over the geodesic neighborhood of that cell. The two encoders share a topology and hold independent weights.

For the evaluation set of 100 structures, we evaluated on 768 simulated particles per structure at the signal-to-noise 0.02–0.30 on the order-3 grid. ARCHER reaches a median angular error of 5.0° and places 77% of particles within 15°. It selects the exactly correct grid cell, one of 36,864, for 41% of particles (shown in Fig. 5).

Moreover, transferability of ARCHER to experimental particles holds without retraining. When applied across all five CESPED targets (Fig. 5A), EMPIAR-10280 and 10648 carry point-group symmetry, and rotating such a particle onto a symmetry-related orientation reproduces the same projection image pixel for pixel. Relying on symmetry based on the reference volumes, we minimize the geodesic over the group. The D_2_ target reads initially at 166.8° and after utilizing symmetry-aware, at 9.1°. The correction to D_2_ leaves the three C_1_ targets bit-identical and does not rescue random poses, which stay at 90°. Detecting initial symmetry point-group

On EMPIAR-10076, a mixture of assembly intermediate of the *E. coli* large ribosomal subunit, ARCHER attains a median error of 2.5° with 78% of particles within 5° of the deposited pose (Fig. S1). The learned classifier’s grid argmax gives 6.16°. Re-scoring against the reference with a shell-normalized matched filter gives 4.53°, and the curvature step gives 2.54°. Since the refinement step searches the whole grid, the ground-truth precision is reached by the classical stage (Fig. 5A).

We ran cryoPARES^28^ on the four targets by first fitting to each of the targets and scored on the particles it did not train on (Fig. 5A and Table S13). Median error across them is 9.4 for the network, 7.5 for the full pipeline, 6.5 classical and 4.2° for cryoPARES (Fig. 5A). It is the more accurate estimator wherever it can be applied, and that qualification is the whole of the difference between the two methods. The precision did not stay the same, once we reduced the fraction of particles placed within 15° for fitting. Here, the classical method leads at 0.818, followed by cryoPARES at 0.805 and our pipeline at 0.718. cryoPARES is more precise when it fits on abundance of particles per structure. Median and tail measure different things here, and we report both in Table (S13).

Since cryoPARES is fit to one specimen at a time, we measured the performance on untrained targets. Applying each of the four trained models to each of the four targets, on the same particles and with the network stage alone, gives a median of 3.9° to 8.8° on the diagonal and 80° to 135° everywhere else (see Table S16). The off-diagonal values sit at the chance level of each target — 120° at C_1_, 97° at C_2_, 90° at D_2_. The comparison in Fig. 5A displays the model excellent on the single structure it is fit to, but not transferable to the others.

Reference-free estimators solve a harder problem – no reference, no translations – and are compared separately in the supplementary Fig. (S3) rather than alongside here. We modulated input data size for CryoFastAR on a homogeneous target in the supplementary Fig. (S2). They are not weak baselines when given enough data. cryoDRGN’s ab-initio schedule sits at chance on the 8,000 particles these arms are scored on, but at 100,000 reaches 1.96, 5.32 and 17.09° on the three C_1_ targets, ahead of our network on each; it degrades on the two symmetric targets, for which its homogeneous entry point offers no symmetry option. Their for these reference-free models incur more inference cost and risks poor transferability.

Errors concentrate in a band of viewing directions rather than spreading over the sphere (Fig. 6B). The residual error of ARCHER is localized in orientation rather than diffusive, which is what makes it addressable by better candidate selection. Furthermore, an uneven distribution of viewing directions attenuates resolution on its own, through a sampling compensation factor that weights each direction by how often it is occupied^88^.

### B. Reconstructions quality

Pose accuracy matters mainly through the map it produces. We evaluate with the CESPED benchmark protocol^31^, which reconstructs each half-set independently at the predicted orientations and scores the result against the deposited structure.

On EMPIAR-10409, ARCHER reconstructs to 3.634 Å against the deposited map. The classical matched filter reaches 3.625 Å and cryoPARES^28^ 3.477 Å: all three lie within 0.16 Å of one another, and their half-map resolutions span 0.05 Å (Fig. 5C). On this measure ARCHER is comparable in reconstruction quality to the supervised method.

The scoring is not uniform across the benchmark. Scored the same way on all four targets, our reconstructions trail cryoPARES by 0.16 Å and 0.22 Å on EMPIAR-10409 and 10648 but by 2.11 Å and 2.06 Å on EMPIAR-10166 and 10280 (Table S15). Feeding more particles improved our reconstruction quality. We reached cryoPARES’s quality when supplementing 203,000 and 117,478 on EMPIAR-10409 and EMPIAR-10648.

Now, we questioned whether the resolution of reconstruction could reach any more meaningful levels. On EMPIAR-10409 the masked shell correlation never falls to 0.143 within the band (Fig. 11C). All reconstructions of classical, cryoPARES, and ARCHER are already at the 2.94 Å Nyquist of the native sampling for EMPIAR-10409 and 2.99 Å for EMPIAR-10648 (Table S15). Therefore, a small difference in the resolution score does not guarantee a better map. Fig. (11) shows how far that score moves under masking alone.

On five CESPED targets spanning box sizes from 136 to 284 voxels and both 200 and 300 kV optics, ARCHER reproduces the deposited density on every one of them (Fig. 3A). Embedded in a common principal-component space, the five targets occupy distinguishable regions (Fig. 3B). Although data collection differs in defocus, contrast and sampling, the molecules in feature space are relatively close. Measured instead against the Nyquist limit of the 64^3^ working box the estimate is made in, the reconstructions sit within a factor of 1.0–1.4 of that limit on all five (Fig. 10). The map quality reported above for EMPIAR-10409 (3.63 Å) is measured at the native 1.47 Å sampling.

**FIG. 3.**
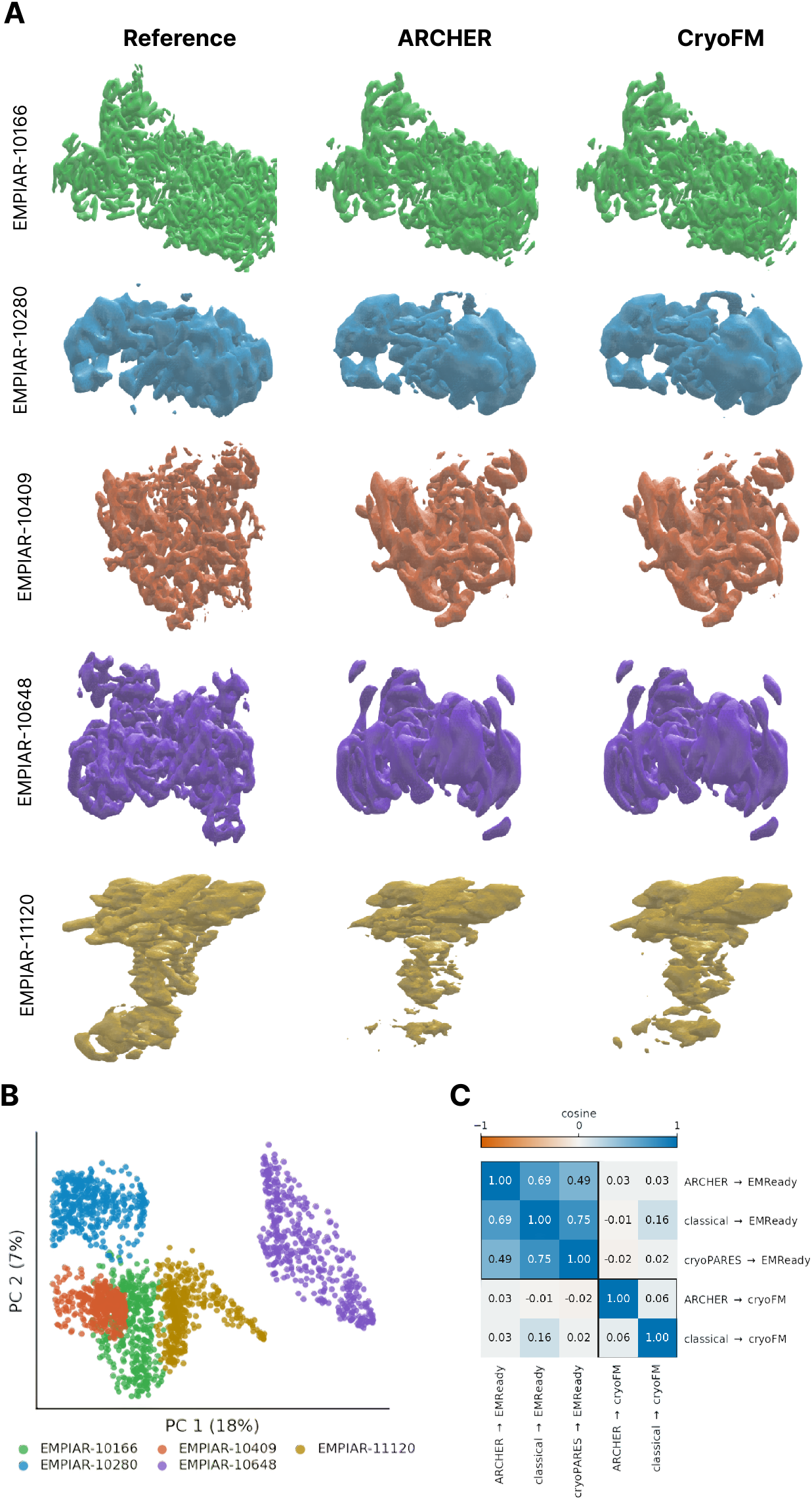
Reconstructions and restorations across specimen. (A) Isosurfaces for four CESPED targets, one row per target and one column per map: the deposited reference, the ARCHER reconstruction, and that reconstruction after cryoFM restoration^36^. Color identifies the specimen; levels enclose a fixed fraction of the mask volume within each row, that is, among maps of the same molecule. (B) Every point is one experimental particle image embedded by principal component analysis, colored by dataset. The four specimens occupy distinguishable regions, and the separation is structural rather than instrumental: repeating the embedding on phase-scrambled particles, which preserves each image’s power spectrum while destroying its structure, collapses the between-specimen separation from 1.06 to 0.05 in units of the within-specimen spread. (C) Cosines between restoration displacement vectors in full voxel space. A restorer moves every map it is given in nearly the same direction, whichever estimator produced that map, so restoration acts on the map rather than on the pose estimate that made it. The same reconstructions embedded by method are Fig. S4.

**FIG. 4.**
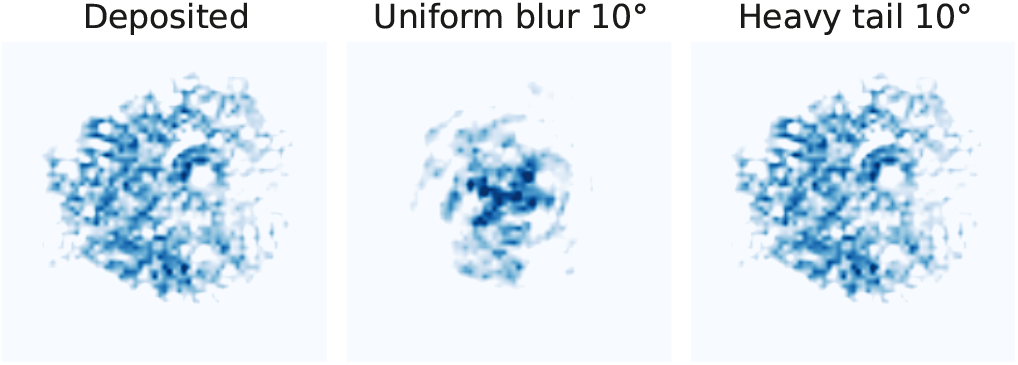
Error shape governs the map. Reconstructions at matched mean angular error under uniform and heavy-tailed pose perturbation.

#### a. Error shape governs the map

What limits a reconstruction is not the average pose error but its shape. Perturbing the deposited poses by a uniform 10° blur degrades the map to 6.5 Å, while a heavy-tailed perturbation of the same mean leaves it at the 2.94 Å Nyquist limit; at a mean of 20° the two differ by more than 11 Å (Figs. 6C and 4). The pose estimators can tolerate a few catastrophically wrong orientations and average away but uniform errors on every particle distort reconstruction. cryoPARES carries the largest mean angular error measured here, 41.3°, and still produces good maps at 3.48 Å, better than a uniform 10° blur, while ARCHER and the classical filter sit on top of one another at 22° and 3.63 Å.

#### b. Restoration as a downstream operator

Here we test whether ARCHER is compatible with a learned 3D cryo-EM restorer, which benefits downstream refinement. EMReady^39^ raises the shell-correlation area from 0.674 to 0.697 and the masked correlation to 0.809 (Fig. 7), and cryoFM^36^ sharpens the grid-argmax reconstruction, before refinement, from 4.10 Å to 3.68 Å. On strongly shuffled poses, both restorers suffer on the resulting low-quality reconstructions. Furthermore, the two restorers do not behave the same way when combined with different pose estimators (Fig. 3C): EMReady moves three separate estimated maps in nearly the same direction, mean cosine 0.64, while cryoFM’s displacements are near-orthogonal at 0.06. On the pose-estimator’s manifold (supplementary Fig. S4), maps from different pose estimators fall close together compared to the distorted maps.

Overall, ARCHER works straightforward from 2D to 3D – particles pose-estimation and back-projected, while a restorer works merely from 3D to 3D. By processing directly from the data source rather than from a prior over densities, our model potentially eliminates restorer-based biases for final restoration.

### C. Conformational heterogeneity signal

We also show the capability of ARCHER on treating conformational heterogeneity problem. As cryoEM structures are flexible molecules in nature, a pose estimator should capture a diverse population of the poses. We ran RECOVAR^54^ twice on identical particles with identical translations while varying the rotations, and compared the two per-particle embeddings by canonical correlation^89^. Canonical correlation pairs each axis of one embedding with the axis of the other it best matches across particles, and ranks the pairs by how well they agree. Each pair is one collective mode of motion, and the ranking is blind to the arbitrary linear map a latent space is defined up to.

On EMPIAR-10076 the leading correlation between the embedding computed from our poses and the one computed from the deposited poses is 0.97, and it stays consistent from two to twenty latent dimensions while the mean over all directions falls (Fig. 6A). What the declining mean measures is where extra canonical directions stop carrying conformational signal and start carrying noise, so the motion our poses preserve is concentrated in the few canonical directions.

On EMPIAR-10409, where our angular error is an order of magnitude larger, our poses retain 0.323 of the embedding against 0.311 for the classical matched filter and 0.248 for cryoPARES, with disjoint bootstrap intervals and a floor of 0.02 from the same pipeline run on the same particles with their poses shuffled (Fig. 5B). Here, ARCHER leads other estimators with comparable angular errors above; cryoPARES holds the best median angular error on this target (Sec. IV A) but keeps the least conformational signal, 23% less than our poses. Being accurate about one consensus structure does not guarantee the ability to capture the dynamics around it, and thus, ARCHER is well-suited to generalize across specimens while preserving heterogeneity signal.

**FIG. 5.**
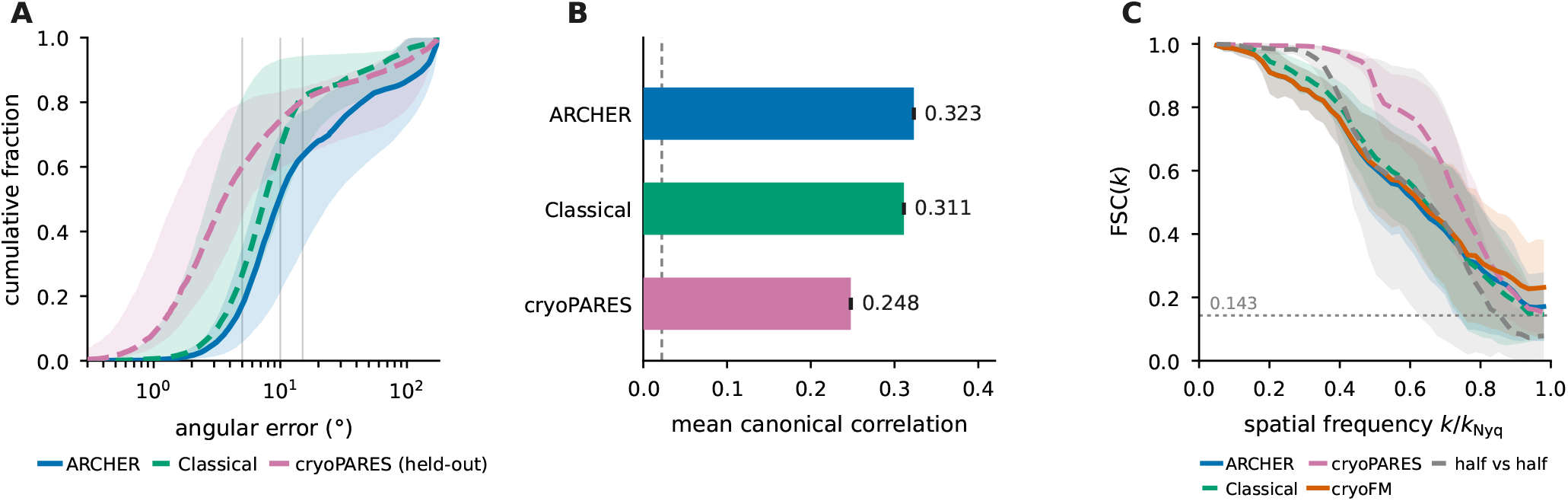
ARCHER performance on benchmark datasets used by per-structure estimators. (A) Cumulative angular error over the five CESPED targets; lines are the median across specimens, shading their full range. (B) Retained conformational signal on EMPIAR-10409, as the mean canonical correlation over the four latent directions; the dashed line is the shuffled-pose floor. (C) Shell correlation pooled over the four targets of Fig. 3A, inside the benchmark mask; the abscissa is a fraction of each target’s own Nyquist frequency and the dashed line marks the 0.143 criterion^18^. Scoring conventions, matched particle counts and error bars are given in App. A.

**FIG. 6.**
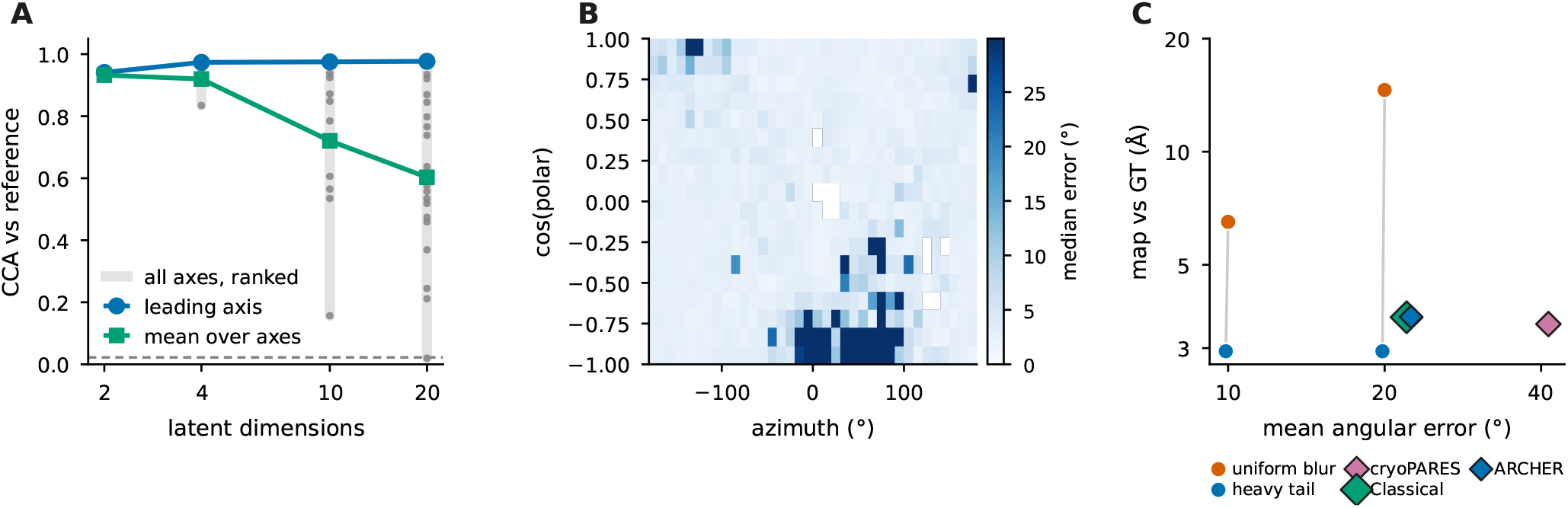
Error quantification from reconstruction maps. (A) Canonical correlation against latent dimension on EMPIAR-10076: the leading direction is flat while the mean declines, so it is the trailing directions that decorrelate. (B) Median error per cell of the view sphere on EMPIAR-10076. (C) Map resolution against *mean* angular error, log–log. Circles are manufactured perturbations at matched mean and opposite error shape — at a mean of 20° the same particles give 2.94 Å under a heavy tail and 14.6 Å under a uniform blur — and diamonds are the real estimators, which sit far to the right of those controls and far below them.

**FIG. 7.**
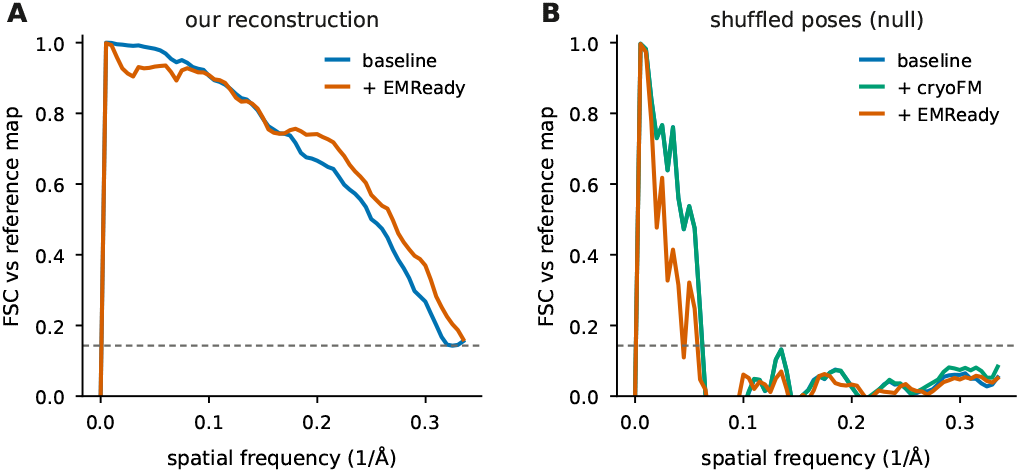
Map restoration. Shell correlation before and after restoration.

**FIG. 8.**
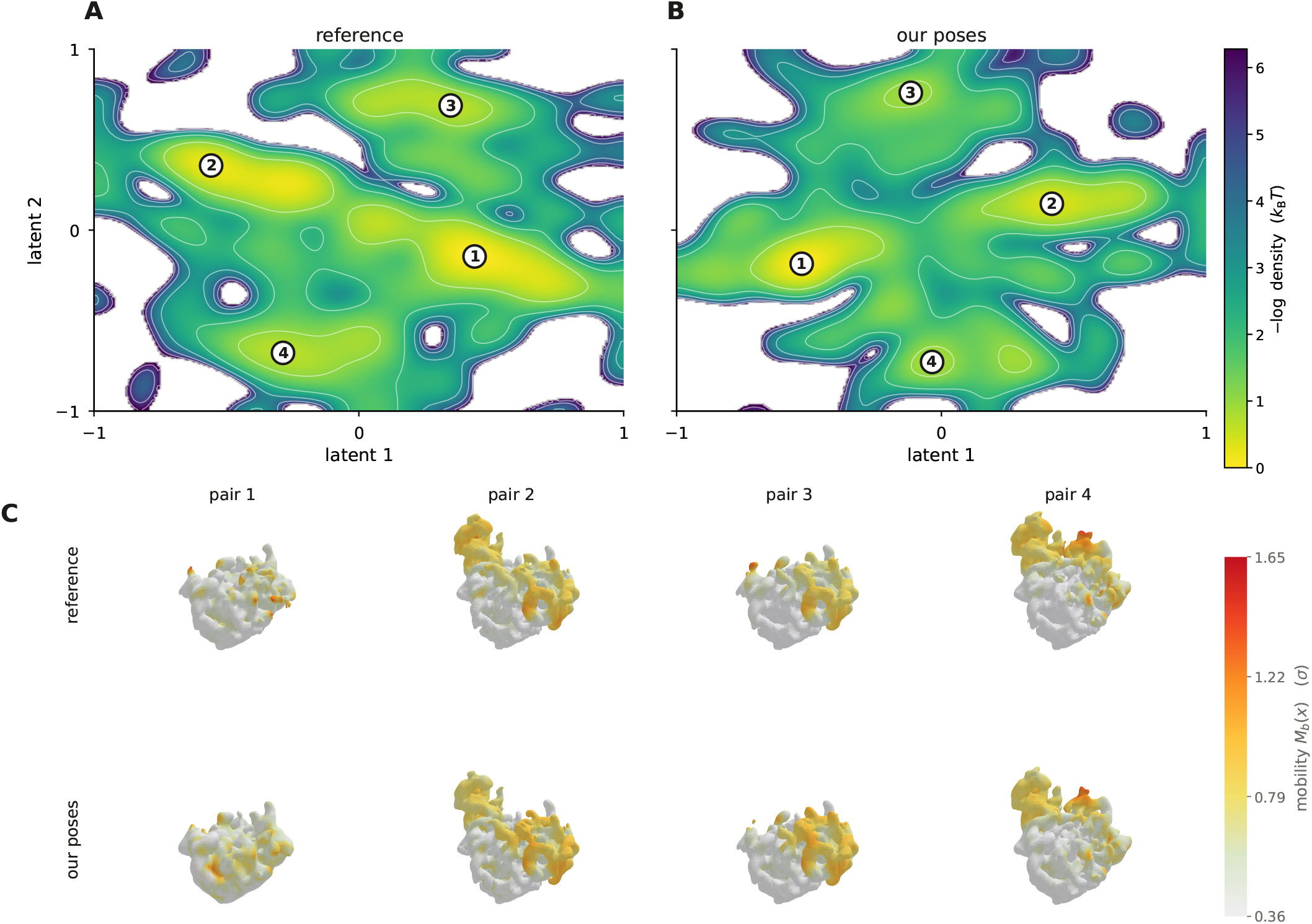
Conformational landscape. (A, B) Deconvolved conformational density over the leading two latent dimensions for the same 91,899 EMPIAR-10076 particles embedded twice, with the deposited poses and with ours, plotted as − log *p* in units of *k*B*T*. Both panels use the same deconvolution weight, and each latent axis is normalized by its own bounds; markers are basins. (C) The reconstruction at each basin, colored by the mobility of Eq. (18) in units of the map’s own standard deviation. Columns pair basins by map correlation: the two latent bases are independent, so basin *i* of one panel need not be basin *i* of the other.

The conformational landscape survives as well as the individual poses. Fig. (8) plots RECOVAR’s deconvolved density for the same particles embedded twice, with the deposited poses and with ours. Matching basins between the two pose sets by map correlation gives 0.92–0.99, and their mobility fields agree at 0.90–0.94. ARCHER’s poses therefore reproduce not only the average structure but where the molecule is localized.

### D. Information scaling on orientation and transfer learning

Amortization across proteins is workable to the extent that something invariant is learnable. A natural candidate is the way orientation information is distributed over spatial frequency, which follows from the central-slice theorem and the microscope rather than from the molecule.

#### a. Orientation information grows as k^2^

Eq. (7) predicts an exponent of exactly 2 with no free parameter. Measured on real particles at their deposited orientations, across models that differ in input channels, training length and box size, the exponent is 1.89 with an autocorrelation-robust standard error of 0.042 (Fig. 9A). The scaling that makes one estimator viable across proteins is obeyed by the data itself, with no network involved, letting an amortized model like ARCHER processing structures that were never trained on.

**FIG. 9.**
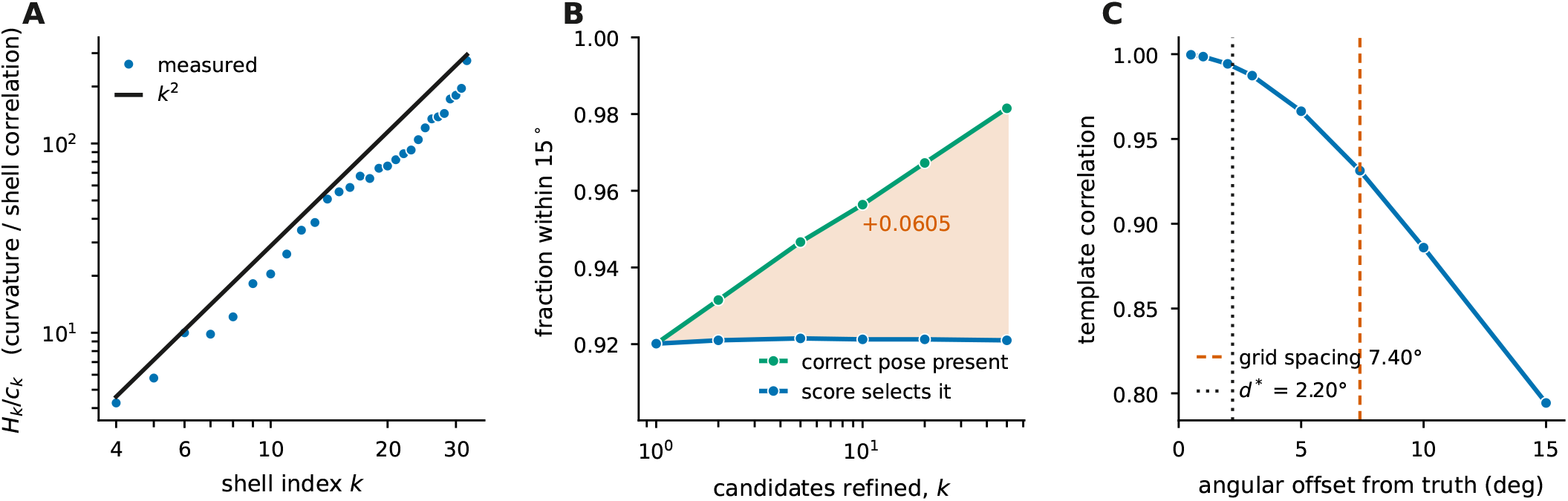
Information, selection and scale. (A) Per-shell curvature normalized by the correlation *c*_*k*_ that shell attains at the true pose, against shell radius, with the *k*^2^ prediction of Eq. (6). The measured exponent over *k* = 4–32 is 1.89, with an ordinary least-squares standard error of 0.037 and an autocorrelation-robust one of 0.042; the latter is the one to read, since neighboring shells are correlated. (B) Fraction of particles for which a correct pose is present among refined candidates, against the fraction for which the score selects it. (C) The score landscape around the true pose, showing the grid spacing and the curvature scale.

#### b. Where the remaining error lives

The error that remains is a selection problem. Refining the top candidates for each particle puts a pose within 15° of the truth for 98% of particles, while the matched-filter score picks that pose for only 92% (Fig. 9B). The right answer is usually already in the candidate set and the ranking rule discards it, so the improvement available to ARCHER is in scoring rather than in a finer grid or a larger network.

#### c. Two angular scales

The score landscape around the true pose sets the two scales the pipeline works between (Fig. 9C). The grid spacing of 7.4° fixes what a discrete search can resolve, and the curvature of the score fixes what a continuous step can add below it. A derivative-free Newton step on three body axes carries the estimate from 4.5° to 2.5° on EMPIAR-10076, a 44% reduction in median error and the largest single gain in the pipeline (supplementary Fig. S1).

#### d. Sampling and measurement limits

Here, we look deeper into the two limits that are much closer across pose estimators than just comparing the pose error: the resolution the working box can support, and the precision with which shell correlation can tell two estimators apart. First, the pose estimate is done on an *L* = 64 crop, and no pose can carry a reconstruction past that crop’s Nyquist frequency. Across the five CESPED targets the reconstructions sit within a factor of 1.00 to 1.39 of it, and a consensus reconstruction from the deposited poses at the same box reaches the same limit, so further pose accuracy would not be meaningful (Fig. 10).

**FIG. 10.**
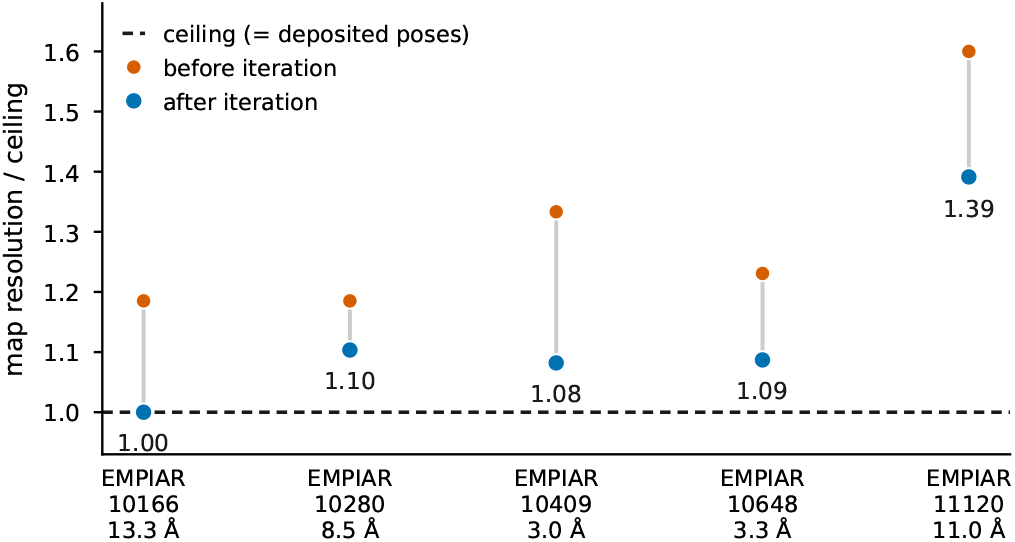
Cross-target reconstruction against the sampling bound. Five CESPED targets (EMPIAR-10166, 10280, 10409, 10648, 11120), each evaluated against the Nyquist limit of of the 64^3^ box the pose estimate is made in. The y-axis is the ratio of map resolution to that ceiling, before and after hierarchical refinement.

The second limit is the measurement itself (Fig. 11). A fixed 0.143 threshold holds every shell to a standard the innermost ones cannot meet, since a shell at radius *k* holds only about 4*πk*^2^ samples, which is what the 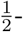 bit criterion corrects by scaling the threshold with that count^19^, and a solvent mask (applied before the transform) raises the correlation while making it less certain – where the inflation high-resolution noise substitution was introduced to detect^90^. Scored both ways on the same reconstruction the reported resolution moves from 3.57 Å unmasked to 2.94 Å masked, by at least 0.63 Å since the masked curve never crosses 0.143 within the band, while the three pose estimators scored through that same routine span 0.12 Å.^91^ The choice of mask moves the headline number five times further than the choice of method does, which is the sense in which these estimators are not separated by resolution. Separating them needs a quantity beyond angular error, and retained conformational signal is the one this work finds discriminating.

**FIG. 11.**
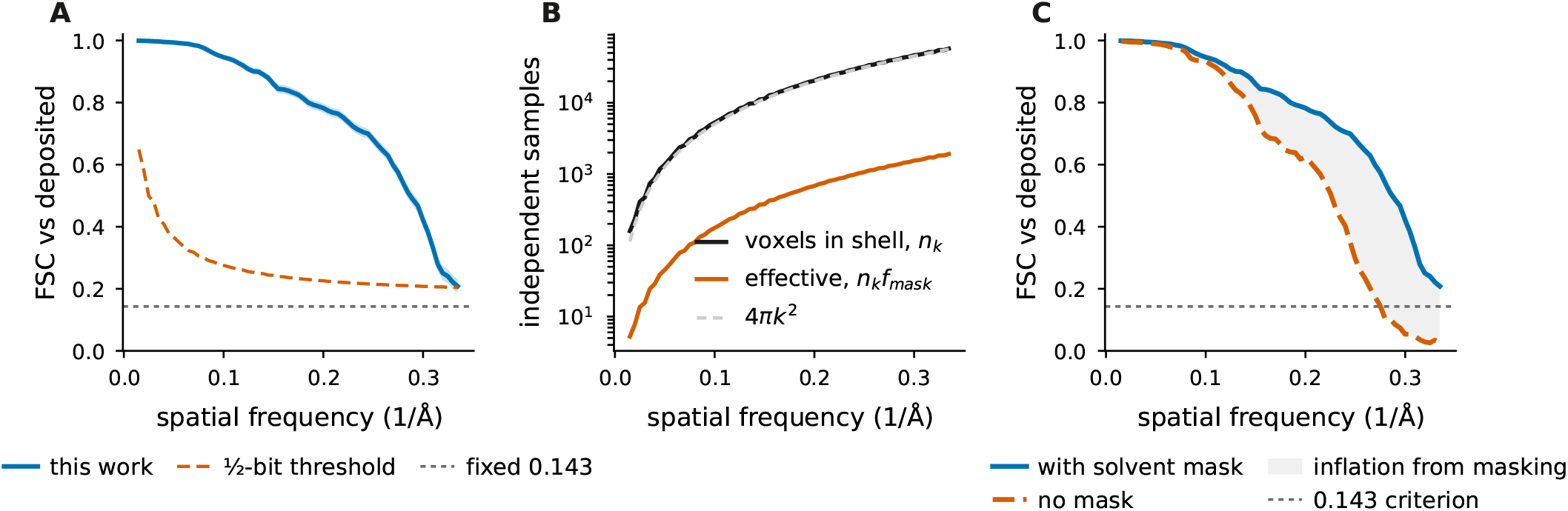
Shell correlation carries its own statistics. All three panels are the ARCHER reconstruction of EMPIAR-10409 scored against the deposited map, at the native 1.47 Å sampling. (A) The reconstruction against the fixed 0.143 criterion and against the 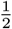-bit criterion, which scales with the number of samples in the shell^19^; the two disagree most where samples are fewest. (B) Samples per shell, following 4*πk*^2^, and the effective count after masking, which is lower by the mask’s volume fraction (3.3% here). (C) The same reconstruction scored masked and unmasked: masking raises the curve throughout, and here pushes the 0.143 crossing past Nyquist altogether, so the shift in reported resolution is a lower bound of 0.63 Å rather than a measured value. The classical arm and cryoPARES are censored at the same 2.94 Å, so on this target the masked criterion cannot separate the three.

## V. DISCUSSION

Conditioning a pose estimator on the reference volume moves the specimen out of the weights and into the input. The consequence is that one set of parameters serves proteins that were never in training, at accuracy competitive with methods fit to each dataset.

The argument has both theoretical and empirical support. Bayes’ rule applied to Eq. (2) makes *p*(**R** | *y, V*) a single functional of the transfer function, the noise spectrum and the slice geometry, with the specimen entering multiplicatively through *P*_*k*_ and *R*_*g*_. The likelihood is not specimen-dependent, and a per-structure method obeys the same Eq. (6) that we do – so the frequency scaling explains what a pose estimator must be sensitive to, not why amortizing across proteins should work.

We have presented empirical results on amortization. The first is that a finite-capacity network approximates that functional uniformly enough over the distribution of references to be useful on proteins outside its training set, while the second being that pooling specimens buys more in variance than it costs in the bias of sharing capacity between them. Neither follows from Eq. (6). Both are what the held-out and cross-target results measure, and they are also why a per-specimen estimator such as cryoPARES retains an advantage in angular error on the dataset it was fitted to. Thus, our shared network across specimen shows advantageous and transferable performance across targets.

This clarifies the relationship to existing families. Per-dataset amortized methods^21,23^ learn an encoder and a volume jointly, which is the right design when no reference exists and the goal is ab-initio structure. Supervised per-dataset estimators^28^ learn the same matching operator we do but tie it to one molecule. ARCHER occupies the regime where an initial reference is available, e.g., a consensus map, a homologue, or a predicted structure^42–44^, and the task is to assign orientations at scale. The three regimes are complementary, and the present results suggest the shared component is larger than the protein-specific one.

Although per-structure models indistinguishably obtain high resolution in reconstruction (Table S15), what separate them is how much conformational signal survives to the downstream analysis, and that is the measure on which our poses lead every estimator compared here – 0.323 of the embedding retained against 0.311 and 0.248, with the most angularly accurate estimator retaining the least. For flexible assemblies that is the quantity of interest, and it should be reported alongside resolution.

Two limitations bound the scope. The estimator requires a reference, and the measurements locate that requirement precisely: it is needed to initialize the loop rather than to sustain it. After the first pass, every subsequent round scores particles against a volume rebuilt from our own estimates, and the accuracy attained with that self-generated reference matches the one attained against an independent reference built from the opposite half-set (4.54° against 4.53° at the grid stage, and 2.60° against 2.54° after refinement), holding to within 0.1° over three further rounds. Starting the same loop from a deliberately crude reference behaves differently: accuracy falls as the iterations proceed, because a poor reference yields a poor reconstruction and the error compounds. The reference must therefore be good enough to place the first estimate in the right basin – a consensus map, a homologue, or a predicted structure – after which the pipeline supplies its own. And the residual error is a selection problem: a correct pose sits in the candidate set for 98% of particles while the score picks it for 92%, so a better ranking rule is the clearest available gain. A learned selector over refined candidates is the natural next step, and the measured headroom quantifies what it can deliver.

## VI. CONCLUSION

Orientation assignments (and other problems) in cry-oEM have been treated as a per-dataset problem because the reference volume has always been carried in the parameters of whatever performs the assignment. In this work, we introduced ARCHER as a general solution for this problem, where it can be applied for structures (seen or unseen by the model).

Notably, the main advantage here is not accuracy but rather generality. ARCHER can be applied to various structures, but its cost is visible. Specifically, when a target’s sampling does not bind, we trail behind a specimen-trained estimator by about 2 Å. When sampling does bind, angular error stops discriminating among competent estimators, and the measurement that continues to separate them is how much conformational signal reaches the downstream analysis. We propose reporting that alongside resolution whenever flexible assemblies are the object of study.

Overall, the performance shown by ARCHER is not trivial. It is able to reconstruct unseen structures, using conditionals obtained from reference structures, within competitive error margins – a notable feature for a rather general technique with respect to the prior mentioned related works.

## ACKNOWLEDGMENTS

The authors declare no competing interests.

## VII. CODE AVAILABILITY

Codes and data are deposited under https://github.com/ndnng/ARCHER

## Supplementary Material

for “ARCHER: Amortized cross-specimen pose estimation for cryo-electron microscopy” Nhan D. Nguyen and Bao Pham

**TABLE S1.**
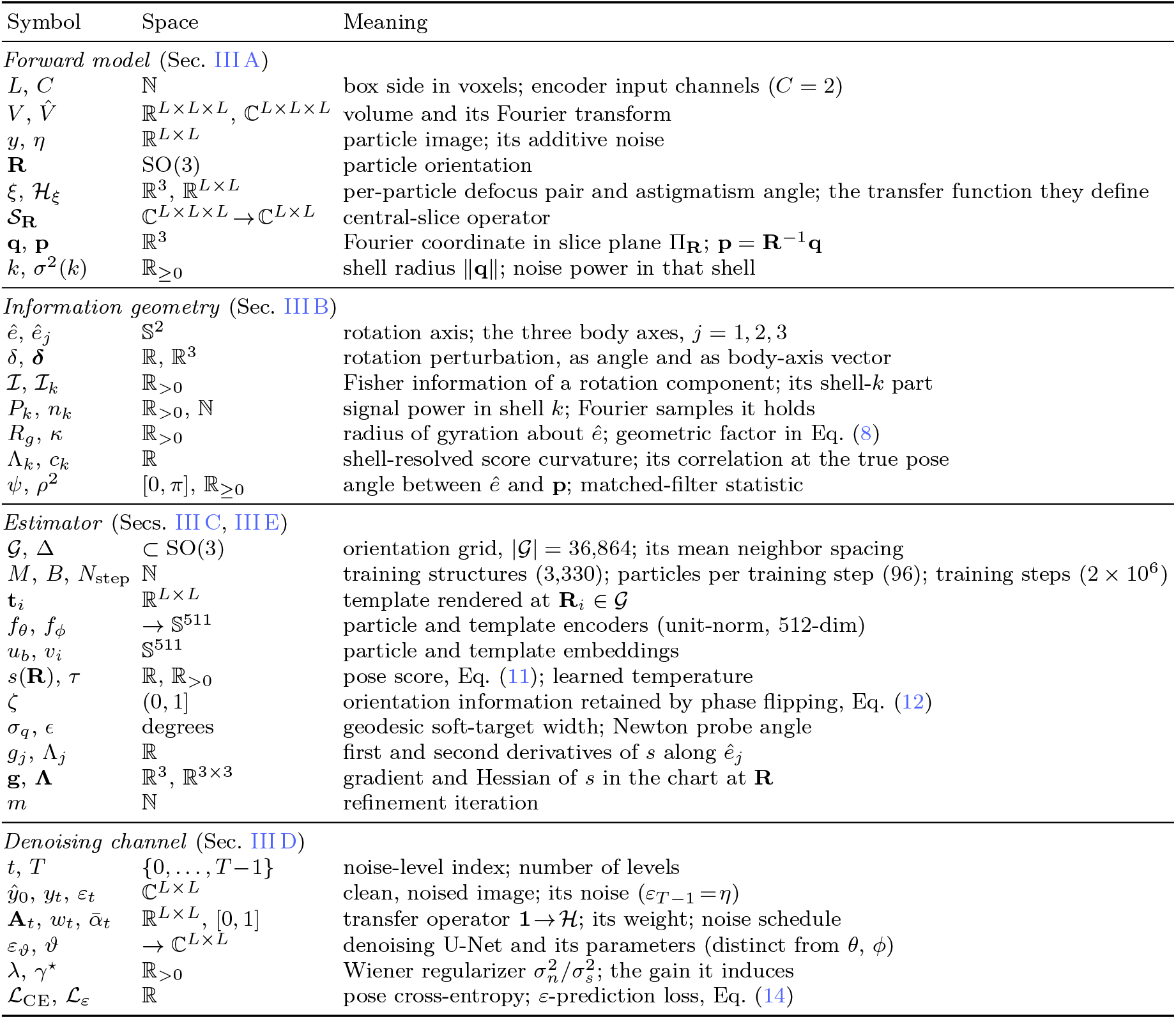
Notation. Each symbol carries one meaning throughout; *L* is reserved for the box side, *C* for a channel count, and *H* for the contrast transfer function, following the convention of Refs.^8,67^.

## Appendix A Benchmark scoring details

### Matched particles (Fig. 5A)

Each estimator is scored on data it did not train on. Our encoder and the classical filter never see any of these proteins, so all 8,000 particles per target are held out for both; cryoPARES, being trained per specimen, is scored on the subset lying outside its own training split. Restricting our two arms to that same subset moves their medians by at most 0.09°, so the curves stay comparable.

### Error bars (Fig. 5B)

Bootstrap over particles, 200 resamples.

### Shell correlation (Fig. 5C)

Every arm is reconstructed from the same number of particles per half-set. The band is one standard deviation across targets. The method curves are measured against the deposited map, whereas *half vs half* is our own two half-sets against each other.

### Latent spread (Fig. 6A)

Each grey bar spans the strongest to the weakest canonical direction at that dimension and the dots are the individual directions, ranked. The spread is structure, not uncertainty — RECOVAR is deterministic given its inputs — and it shows that the decline in the mean is the trailing directions decorrelating rather than the leading one degrading. The dashed line is the shuffled-pose floor.

### View-sphere coverage (Fig. 6B)

The 20,000 EMPIAR-10076 particles populate 637 of the 648 cells.

### Ordering by mean error (Fig. 6C)

cryoPARES carries the largest mean error of any arm shown while still producing the best map, because its errors are tail-like rather than diffuse.

**FIG. S1.**
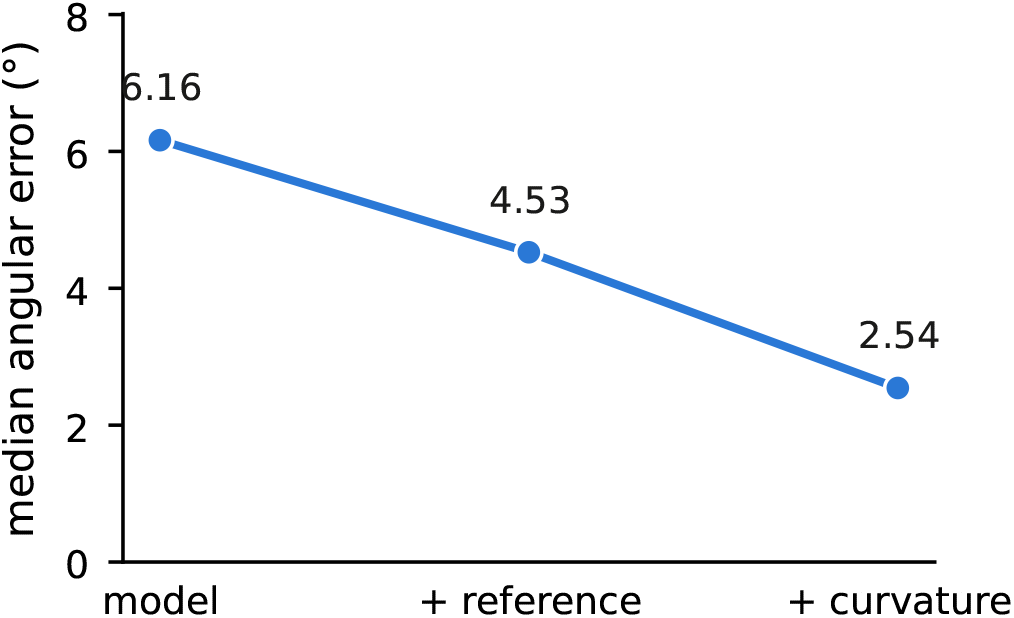
Where the accuracy is gained. Median angular error on EMPIAR-10076 through the refinement ladder: the grid maximum of the learned posterior, re-scoring against a reference rebuilt from those poses, and the curvature step.

**FIG. S2.**
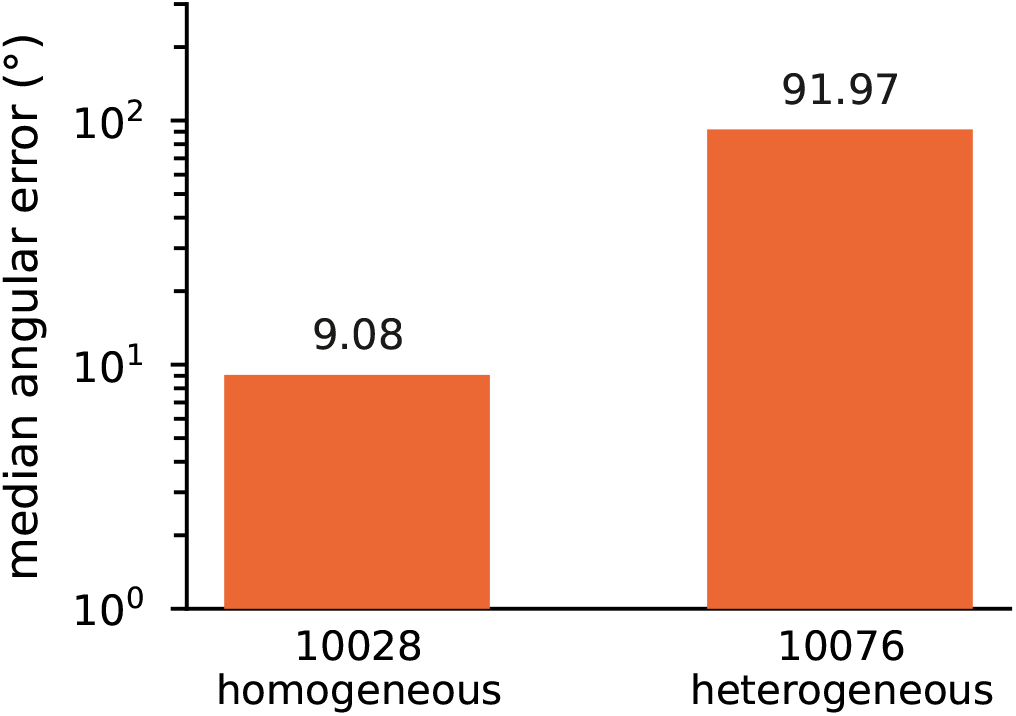
Set-size control for the ab-initio comparison. CryoFastAR evaluated on the homogeneous EMPIAR-10028 at the same particle count used for EMPIAR-10076, isolating specimen heterogeneity from set size.

**FIG. S3.**
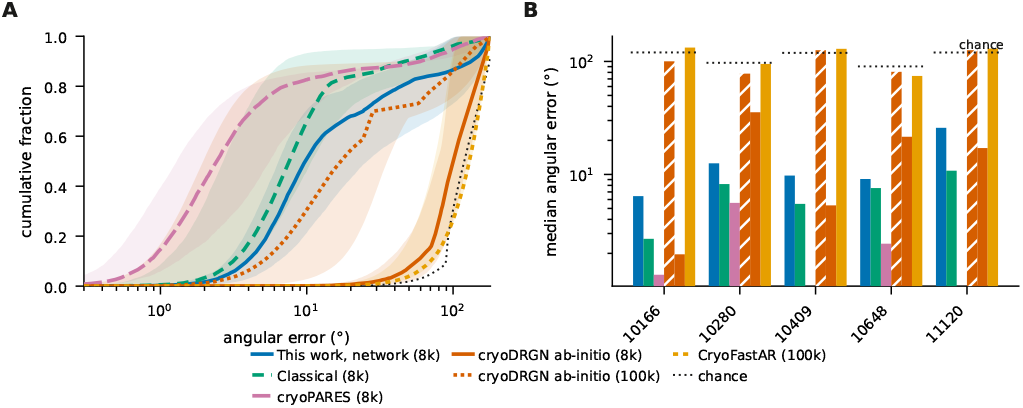
Reference-free and per-specimen baselines on matched particles. (A) Cumulative angular error, median over the five CESPED targets with the full range shaded. (B) Median per target. All arms score the same 8,000 particles per target from one seeded selection; cryoPARES is matched to them by particle identity (`rlnImageName`). Chance is drawn per target, since a point group of order °|*G*| lets a random pose fall nearer an equivalent copy: 120° on the three C1 targets, 97° on the C2 target, 90° on the D2 target. cry-oPARES was trained for four of the five targets and reaches 1.29, 5.59 and 2.44° on the matched particles. EMPIAR-11120 carries no cryoPARES bar because no model exists for it; EMPIAR-10409 carries none in this panel alone, where the name-matched subset was not computed, and its model reaches 5.42° on the unmatched 8,000 (Tables S14, S15, S16). The two reference-free methods solve a harder problem — no reference and no translations — and ran at their default settings, each additionally given a solved global rotation and a choice of handedness beyond the point-group quotient every arm receives. Those allowances can only help them, and a random-pose control through the identical alignment stays at 119.6°, so the alignment manufactures no accuracy. Particle count decides cryoDRGN. At 8,000 particles it sits at chance on every target; at 100,000 it reaches 1.96, 5.32 and 17.09° on the three C1 targets, ahead of our network on all three, breaking from chance only near epoch 14 of 30. It reaches 35.49 (C2) and 21.53° (D2) on the symmetric targets, well above chance but far behind the other arms, as `abinit_homo` exposes no symmetry option. None of these transfers: each is a separate optimization costing 2.6–5.3 GPU-hours. CryoFastAR does not improve with count over the same range.

## Appendix B Architecture and objective

### Encoders

Both encoders in Eq. (11) are residual convolutional networks^73^ with group normalization^92^ and SiLU activations, mapping (*B, C, L, L*) → (*B*, 512) with an *ℓ*_2_-normalized output. They share no weights, and share a topology apart from the stem’s input channels. The forward pass is, in order:

1. **Channel assembly**. The image branch of the reported model takes *C* = 2 channels, [ *y, ℱ* ^−1^[sign(*ℋ*) ℱ*y*]]; the template branch takes *C* = 1. The denoiser of Sec. III D supplies a third channel. Each channel is divided by its own standard deviation, so per-protein contrast does not enter the similarity. Wiener-corrected and whitened channels are implemented but are not part of the reported configuration.
2. **Stem**. One 5 × 5 convolution at stride 2, then SiLU. The stride is not cosmetic: one training step encodes hundreds of templates in one backward pass, and a stride-1 stem would hold every one of them at full 64 × 64 resolution simultaneously, which at the widths used here reaches gigabyte scale for a single activation tensor. Halving the resolution cuts that fourfold and costs little at *L* = 64, where the particle already spans most of the box.
3. **Residual trunk**. Four stride-2 blocks. Each block is `conv3 × 3 → GroupNorm → SiLU → conv3 × 3 → GroupNorm`, added to a shortcut that is the identity when the shape is preserved and a 1 × 1 convolution otherwise, followed by SiLU. Group count is min(8, *C*_out_). Channel width doubles per stage from a base width and is capped at four times that base, so the deepest stages do not dominate the parameter count. For the reported model the base width is 128, giving stage widths [128, 256, 512, 512] and a 2 × 2 trunk output at *L* = 64; the five-stage capacity arm of Table S5 reaches 1 × 1.
4. **Projection head**. Global average pooling to a vector, then `Linear → SiLU → Linear` to dimension 512, then *ℓ*_2_ normalization onto S^511^.

**FIG. S4.**
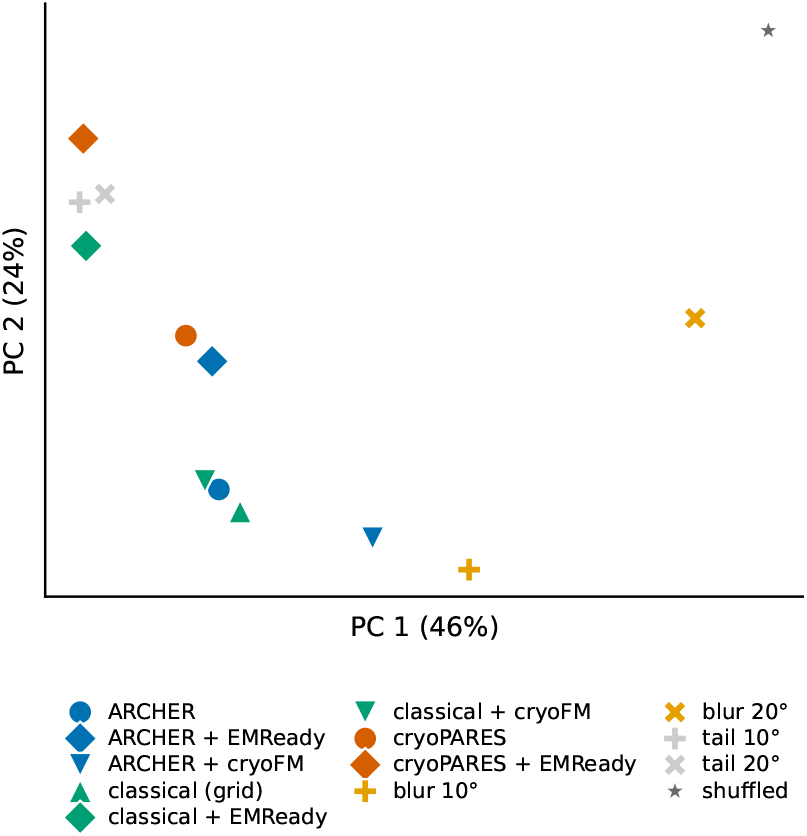
Reconstructions of EMPIAR-10409, by method. Each arm’s map of that one specimen, embedded in the same principal-component space as the projections of Fig. 3B; one point is one map. Restored maps group by restorer rather than by the estimator that produced the input, which is the conclusion the displacement cosines of Fig. 3C reach numerically.

The logit scale *τ* ^−1^ is a single learned scalar, stored as log *τ* ^−1^. Because both embeddings are unit norm, ⟨*u, v*_*k*_⟩ ∈ [− 1, 1], so *τ* ^−1^ is the only quantity setting how peaked the posterior in Eq. (11) can become. It is initialized at *τ* = 0.07 and converges to *τ* = 0.0212 (*τ* ^−1^ = 47.1), a factor of 3.3 sharper than initialization so the scale is genuinely fitted rather than inherited. A clamp at *τ* ^−1^ ≤ 100 guards against an early saturating softmax stalling the gradient; it is never reached.

The reported model has 20.8 M parameters and a 512-dimensional embedding; the capacity ablation of Table S5 uses a 69 M variant differing only in base width, depth and embedding dimension.

#### a. What the architecture deliberately does not contain

There is no volume decoder, no latent variable, no sampling step, no attention and no recurrence. The denoising channel is conditioned on a noise level and evaluated in a single step, so it borrows the training objective of diffusion models without being a generative model; the pose network itself generates nothing: it maps two images to two vectors and the pose posterior is a softmax over their inner products against a *fixed* candidate set. The restriction is deliberate: the volume given poses is a closed-form linear solve (Eq. (15)), so a generative decoder would have to re-learn a projection–slice relationship that is already known exactly.

### 2. CTF handling

The contrast transfer function enters *only* as pixel content, through the phase-flipped second channel *ℱ*^−1^[sign(*ℋ*) *ℱy*], and is never handed to the network as a parameter vector: the network reads the transfer function in the pixels it modulates, which is the same form it takes on a real micrograph. That channel carries information rather than repeating the first — measured on real EMPIAR-10409 particles, the raw and phase-flipped channels correlate at only 0.047.

A Wiener-corrected channel is implemented alongside it, since phase flipping fixes signs but not amplitudes, and it matters most where the CTF has no in-band zero crossing: the *L* = 64 cache Fourier-crops every EMDB map to a fixed box, so large particles land at coarse voxel sizes where the transfer function barely oscillates. It is not part of the reported configuration.

#### a. The denoised channel

The third-channel arm quoted in Sec. III D shares every setting with the reported model and differs only in the added channel, so the comparison of Table S2 is step-matched throughout. The gain is confined to the fine rows: the lead on *<* 15° is at chance, so the coarse assignment is untouched and the channel acts on precision within the correct basin.

**TABLE S2.**
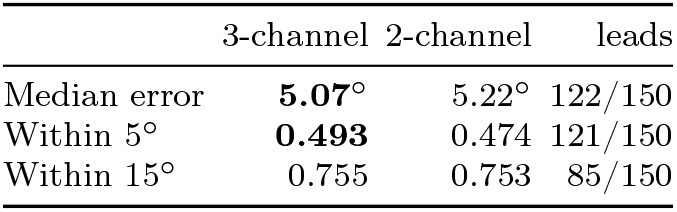
The denoised channel, step-matched. Both arms share every setting and differ only in the added third channel. Means over the last 150 step-matched held-out evaluations, spanning steps 600,000–1,345,000; the lead column counts evaluations at which the 3-channel arm is ahead. Consecutive evaluations of one run are correlated, so those counts describe the trajectory and are not independence tests. The accuracy figures reported in the main text are the 2-channel model.

|  | 3-channel | 2-channel | leads |
| --- | --- | --- | --- |
| Median error | **5.07°** | 5.22° | 122/150 |
| Within 5° | **0.493** | 0.474 | 121/150 |
| Within 15° | 0.755 | 0.753 | 85/150 |

### 3. Objective and optimization

Training minimizes the soft cross-entropy of Eq. (11). The soft-target width is tied to the grid: *σ*_*q*_ = 6° for the reported order-3 model on its 7.40° spacing, and *σ*_*q*_ ≈ = 12° on the 14.78° order-2 grid, i.e. *σ*_*q*_ ≈ 0.81 Δ in both cases. Where a hard target is used instead, label smoothing of is applied, matching the reference SO(3)-classifier implementation.^28,93^

Optimization uses AdamW with decoupled weight decay^94^ of 1 × 10^−5^, held constant across every checkpoint in this work, on a cosine schedule with warm restarts.^95^ The reported model trains at a learning rate of 1 ×10^−4^ with *T*_max_ = 200,000. The short 150k-step order-2 runs of Table S5 use 3 × 10^−4^; that value was inherited by the 69 M capacity arm from its 20.8 M control rather than retuned for it, which is the stated scope limit on that comparison.

#### a. One schedule property worth stating

Setting `T_max` equal to the step budget anneals the learning rate to zero exactly while the model is still improving, so a fixed-budget figure is a lower bound set by that budget rather than a converged value. Reported figures are correspondingly plateau medians over a stated window, and any continuation restarts the schedule.

## Appendix C Full ablation and sweep tables

Throughout this section **bold** marks the best value in a column wherever the column holds directly comparable entries; tables that locate an operating point rather than compare methods carry no emphasis, and the setting adopted is named in the caption.

### 1. Which configuration produced which number

Table S3 states, for every accuracy figure reported anywhere in this work, the grid order, the evaluation noise range, the particle count, the number of proteins and whether point-group symmetry was quotiented. One difference between configurations dominates the rest. The main-text held-out figures are measured on the order-3 grid (36,864 cells, Δ = 7.40°, soft target *σ*_*q*_ = 6°), while the ablation and curriculum tables below (Tables S4–S6 and S12) are sweeps over training choices and were run on the cheaper order-2 grid (4,608 cells, Δ = 14.78°, *σ*_*q*_ = 12°). Halving the grid spacing roughly halves the median error, so the order-2 medians of 10–12° and the order-3 median of 5.0° are one model family measured at two resolutions. Nothing in this work compares an order-2 number against an order-3 one.

### 2. The anneal-endpoint dose–response

Table S4 gives the per-stage detail behind Sec. III A. The hard-noise metric improves monotonically across all five stages, and the legacy metric across the first four before flattening (0.6141 to 0.6130 at the last), so the trend is a dose–response rather than a single point.

**TABLE S3.**
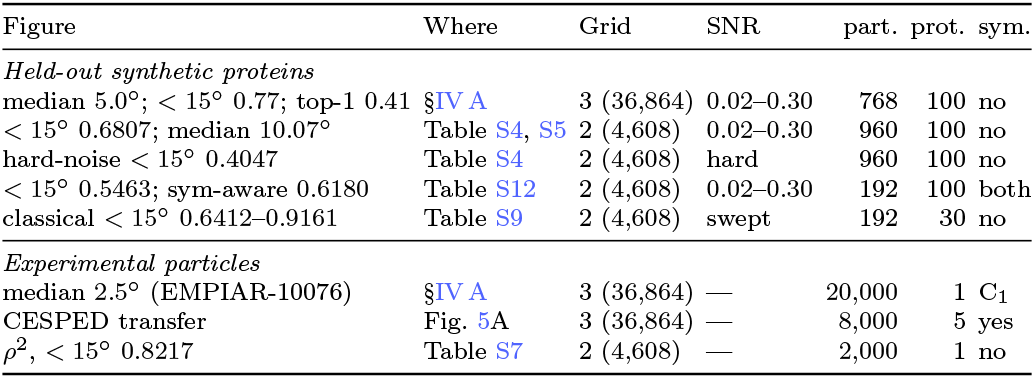
Every reported accuracy figure and the configuration that produced it. *Grid* is the HEALPix order and the resulting cell count; *SNR* is the evaluation noise range, pinned explicitly whenever checkpoints are compared; *sym*. is whether the geodesic is minimized over the point group. The order-2 rows are training-choice sweeps, not the reported model.

**TABLE S4.**
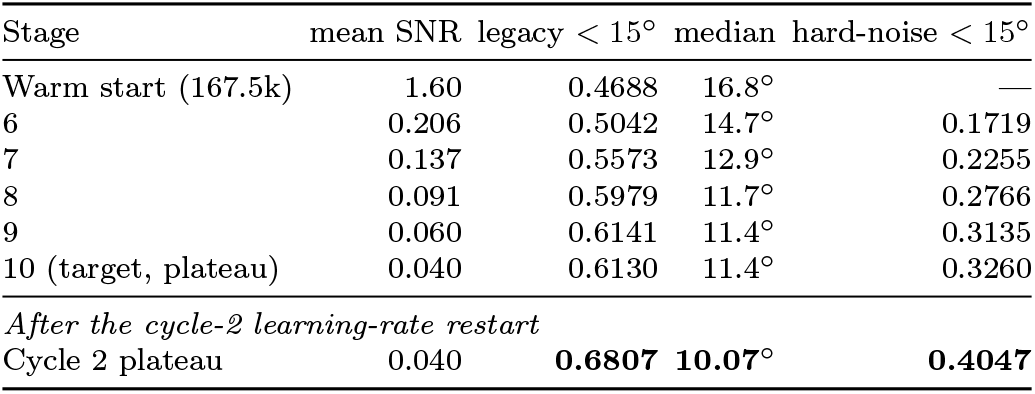
Re-annealing to the calibrated endpoint. The legacy metric is held at the pinned (0.02, 0.30) range in every row, so the rows are directly comparable. Stages are numbered along the geometric anneal (ratio ≈ 0.665 per stage); the warm start enters at stage 6. The stage-10 row is a plateau median over the 12 evaluations of that stage (steps 350k– 377.5k). The cycle-2 row continues the stage-10 arm through a learning-rate restart rather than standing beside it, so it is set below the rule; **that row is the reported model**.

### 3. All three arms at their own plateaus

Table S5 reports the three endpoint arms, each at its own plateau criterion. Table S6 is the controlled version of the same question: there both arms resume the *same* checkpoint under the same schedule and differ only in the SNR bounds.

**TABLE S5.**
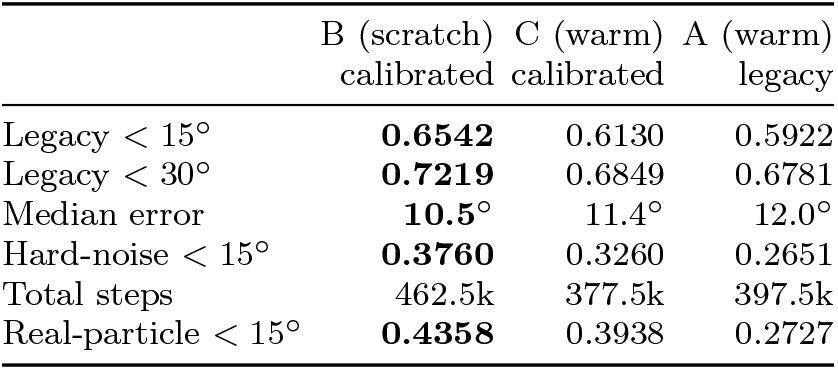
Endpoint arms at their own plateau criteria. Arm B is a from-scratch replication at the calibrated end-point; Arm C is warm-started at the same endpoint; Arm A is warm-started at the legacy endpoint. Bold marks the best of the three. The reported model continues Arm C through a learning-rate restart and exceeds all of them (Table S4, last row).

**TABLE S6.**
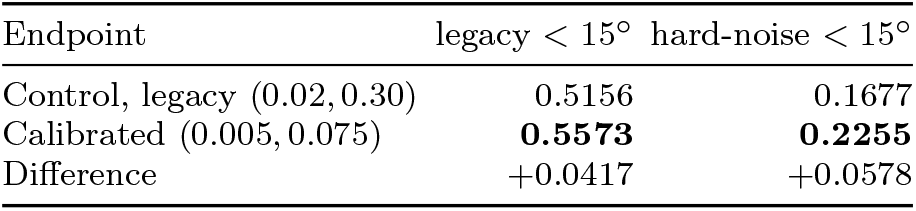
The controlled endpoint comparison. Both arms resume the *same* checkpoint with the same stage count, caps, plateau window, patience, learning-rate schedule and plateau metric; the only difference is the SNR bounds. Medians over the last 10 matched-step evaluations, from 32 matched steps spanning 170k–247.5k. The calibrated arm leads at 32 of 32 matched steps on both metrics. Consecutive evaluations of one run are correlated, so that count is reported as a description of the trajectory and not as an independence test.

### 4. Detectability of real versus synthetic particles

Table S7 is the measurement establishing that experimental particles carry ample pose information at this box size. A single real particle sits 1.7 × above the 2 ln *M* = 16.9 detection threshold and the median particle’s true pose ranks first of 4,608.

**TABLE S7.**
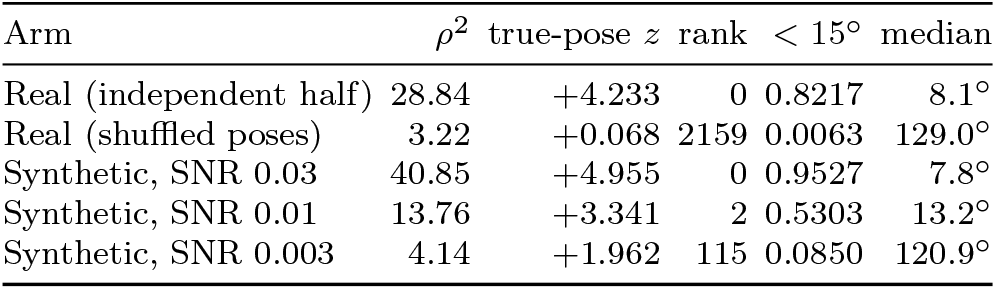
Whitened matched-filter detectability, CESPED EMPIAR-10409 at *L* = 64 (3.128 Å/px, Nyquist 6.26 Å) against a reconstructed reference. *rank* is the rank of the true pose among *M* = 4,608 candidates for the median particle. The real-particle row is the anchor the noise calibration targets; the synthetic rows bracket it. Real particles are `rlnRandomSubset == 2` scored against the half-1 map, so none contributed to the reference; the partly-overlapping variant gives 0.8380 at *ρ*^2^ = 29.44, a 2% difference.

### 5. Rejecting junk particles with the pose confidence

The posterior the pipeline already computes carries a usable quality signal at no extra cost. We score the normalized negative entropy 1+ ∑_*i*_*p*_*i*_ log *p*_*i*_*/* log|*G*|, the same confidence the refinement stage uses, and ask how well it separates real particles from junk. CESPED ships curated particles, so junk is constructed, and the construction is the substance of the test: *phase scramble* randomizes Fourier phases at the measured amplitude spectrum, so every shell carries identical power and only structure distinguishes it; *pure noise* redraws amplitudes from the mean shell profile, giving an image with no particle in it; *off-center* translates a real particle 25–45% of the box, the bad pick that actually populates real datasets. Separation is reported as AUROC, the probability that a real particle outscores a junk one. The reference is built from half 1 and the scored particles are drawn from half 0, so the confidence cannot be reading back its own map.

**TABLE S8.**
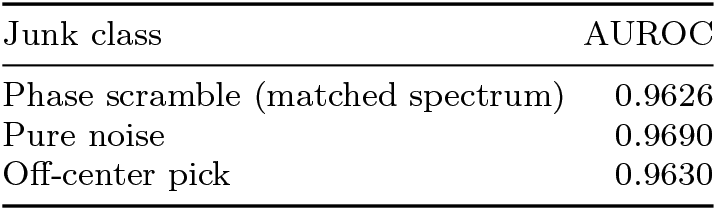
Junk rejection on EMPIAR-10409, 2,000 particles at *L* = 64 against a half-1 reference, for the reported model. AUROC of the pose confidence, real versus each junk class; 0.5 is no separation. Median confidence is 0.364 on real particles against 0.16–0.18 on junk. The phase-scrambled row is the strongest control, since it matches the real amplitude spectrum shell for shell.

The reported model separates all three classes at AU-ROC 0.963–0.969. The phase-scrambled control is the informative one: an image with the real power spectrum and destroyed structure is rejected as readily as pure noise, so the score is responding to particle structure rather than to a difference in image statistics that any variance threshold would catch.

### 6. The behavioral calibration sweep and the symmetry split

Table S9 gives the sweep behind Sec. III A. Averaged over all held-out proteins the classical scorer settles near the ceiling that ambiguity alone implies (Table S10), well short of the 0.8217 it achieves on real particles. Restricted to non-symmetric proteins it reaches 0.8222 at SNR 0.04, matching the real-particle anchor to within 0.0005. The averaged wall is therefore a property of pooling proteins whose poses are not separable, not of the scorer.

**TABLE S9.**
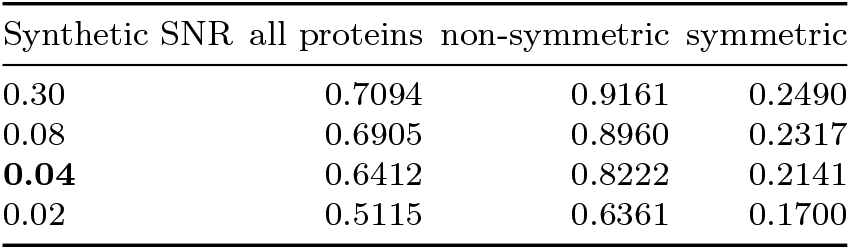
Classical *<* 15° against synthetic SNR, split by whether the protein admits indistinguishable poses. The all-proteins column saturates at the ceiling implied by ambiguity alone; the non-symmetric column crosses the real-data anchor of 0.8217 at SNR 0.04. Bold marks that row, the calibrated endpoint adopted throughout; it is the operating point, not the largest entry.

### 7. Symmetry: the ambiguity audit and its point-group verification

Table S10 gives the ambiguity statistics on the 30-protein sweep set, where the predicted ceiling and the observed classical saturation agree to 0.2%. Table S11 gives the quantization check on the full 100-protein held-out set. Every one of the 26 symmetric proteins lands on an integer point-group order with none scattered in between, which a correlation artefact could not produce; the dominance of C2 and D2/C4 is what an EMDB-derived corpus of oligomeric complexes should look like.

**TABLE S10.**
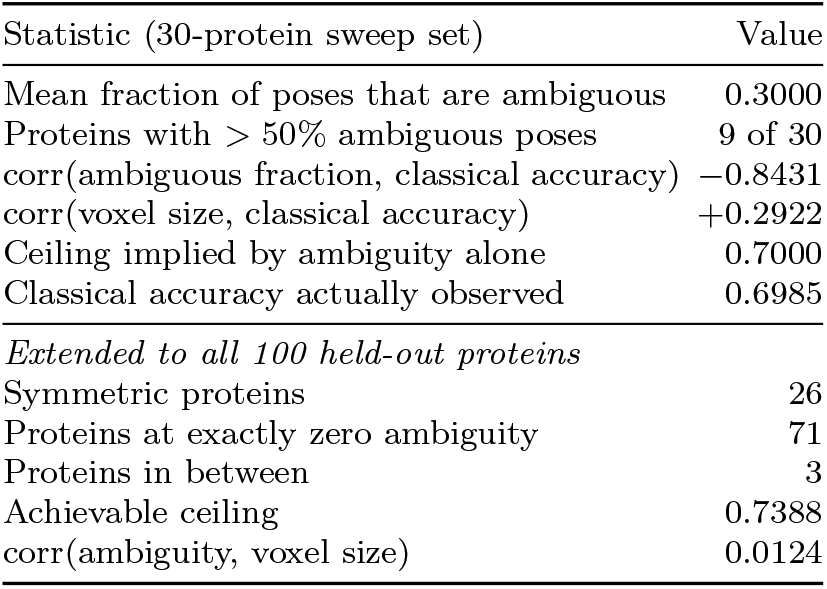
Pose ambiguity measured from the volumes alone — no particles, no noise, no scorer.

**TABLE S11.**
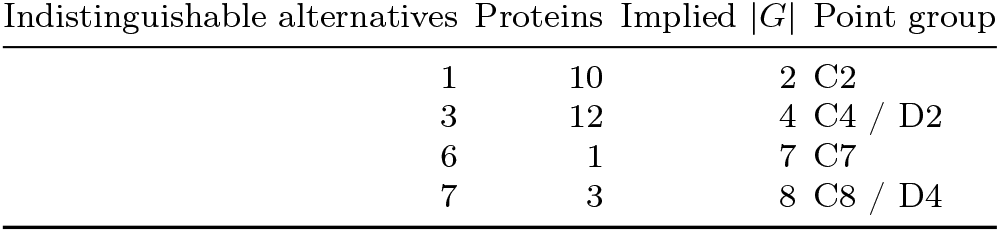
Independent verification of the ambiguity measurement by point-group quantization. A point group of order |*G*| predicts exactly |*G*| − 1 indistinguishable alternatives per pose.

**TABLE S12.**
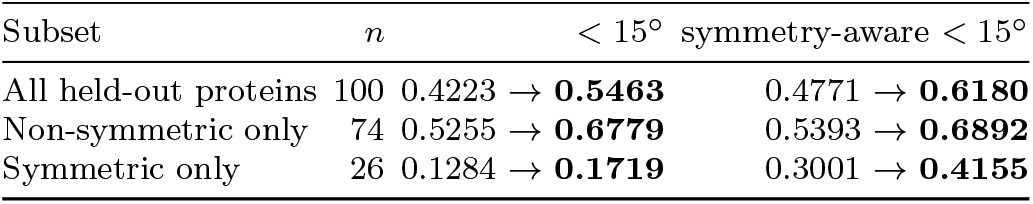
Re-scoring both checkpoints with symmetry separated out. All 100 held-out proteins, 192 particles each, both checkpoints scored at the *same* pinned (0.02, 0.30) range. Arrows run from the 167.5k checkpoint to the plateaued calibrated checkpoint. The non-symmetric row is the control confirming the symmetry-aware correction is not simply inflationary: it moves almost nothing there.

### 8. Per-target accuracy against the per-specimen estimator

Table S13 is the per-target breakdown behind the comparison in Sec. IV A. cryoPARES is scored only where it did not train, and all four arms are scored on exactly those particles, so no arm is credited for particles another arm never saw. Pooling the table by taking the median across the four targets gives medians of 9.5° for the network, 7.5° for our pipeline, 6.6° classical and 4.2° for cryoPARES, and fractions within 15° of 0.689, 0.714, 0.809 and 0.805. The two statistics do not induce the same ordering: cryoPARES leads on the median by a wide margin and ties the classical filter on the tail.

**TABLE S13.**
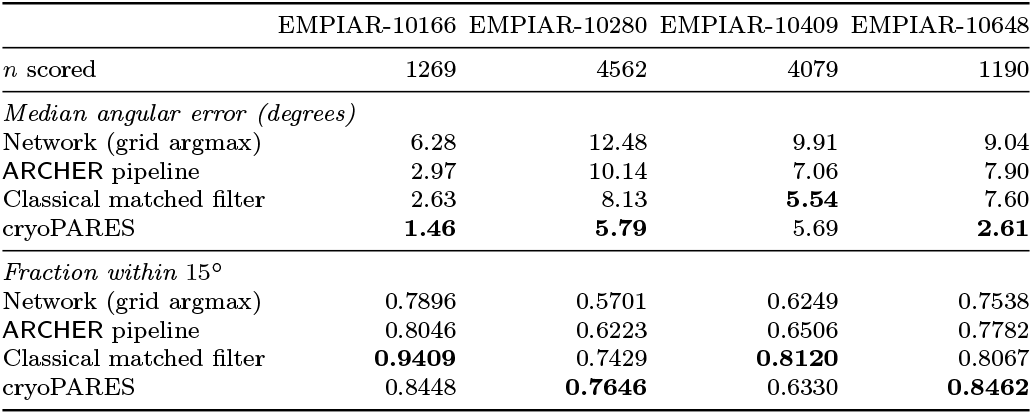
Per-target accuracy on the particles cryoPARES did not train on. All four arms are scored on the *same* particles of each target — the intersection of our seeded 8,000 with the complement of that target’s cryoPARES training split, *n* below — with the same symmetry-aware geodesic, so the columns are directly comparable. Bold marks the best arm per target. cryoPARES holds the best median on three of the four targets while the classical filter holds the largest fraction within 15° on two of them: the two statistics rank the arms differently, which is why both are reported. EMPIAR-10409 is the clearest case: cryoPARES leads on neither statistic there, taking a median within 0.15° of the classical filter while placing 0.179 fewer particles within 15° — a tail close to that of our unrefined grid argmax.

**TABLE S14.**
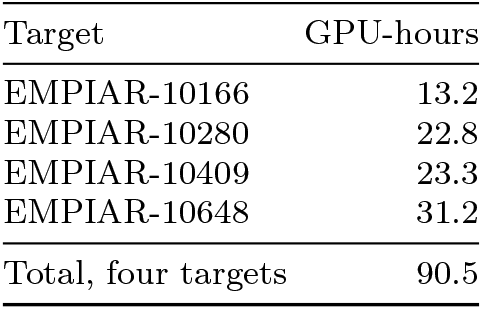
Training cost of the per-specimen comparison. cryoPARES^28^ is supervised on the target it is applied to, so each entry is a separate optimization; times are single-GPU wall clock for the completed run on each dataset at the scope used here (200,000 particles, 40 epochs). ARCHER is trained once across 3,330 structures and applied to every target without retraining, so its cost does not recur per specimen — for a new target it is an inference pass.

| Target | GPU-hours |
| --- | --- |
| EMPIAR-10166 | 13.2 |
| EMPIAR-10280 | 22.8 |
| EMPIAR-10409 | 23.3 |
| EMPIAR-10648 | 31.2 |
| Total, four targets | 90.5 |

## Appendix D Verification of the geometric conventions

Table S17 lists the checks that fix the sign, centering and transfer-function conventions used throughout. Each compares an implementation against an expectation derived outside it — an analytic identity, a reference implementation, or an independent reconstructor — so agreement is evidence rather than self-consistency.

**TABLE S15.**
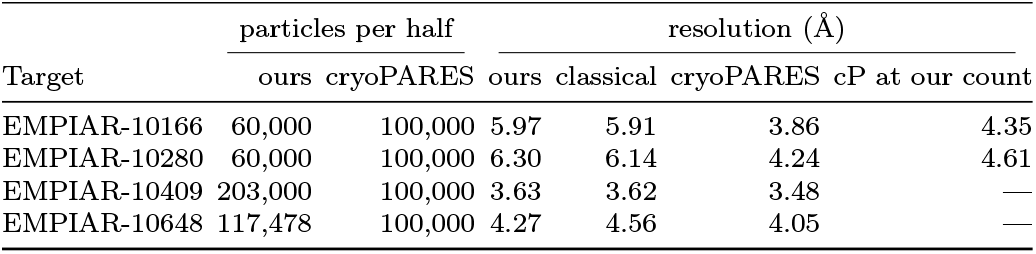
Reconstruction quality per target against the deposited map, as the unmasked 0.143 crossing. Unmasked is reported because the masked curve does not cross before Nyquist on EMPIAR-10409 and 10648, so a masked number there is a floor rather than a measurement (Fig. 11C). Our poses were predicted for 60,000 particles per half on EMPIAR-10166 and 10280 against cryoPARES’s 100,000, so on those two we also rebuilt cryoPARES from the same 60,000 (last column). Matching the count recovers 0.49 and 0.37 Å of the gap and cryoPARES still leads by 1.62 and 1.69 Å, so the deficit there is mostly pose quality and not set size. The reverse control — rebuilding cryoPARES at the *larger* counts we used on EMPIAR-10409 and 10648 — was not run, hence the two dashes; it would widen the gap on those targets rather than narrow it. Entries are rounded to two decimals, which is coarser than the differences discussed in the main text: on EMPIAR-10409 the three values are 3.634 (ours), 3.625 (classical) and 3.477 Å (cryoPARES), a spread of 0.157 Å. The classical arm, reconstructed from the same particles as ours with no learned component, lands within 0.3 Å of our pipeline on every target: the gap to cryoPARES is therefore a property of reference-conditioned refinement at this working resolution and not of the network. On the two targets where we supply *more* particles than cryoPARES the gap is 0.16 Å and 0.22 Å; the former is the agreement quoted in the main text.

**TABLE S16.**
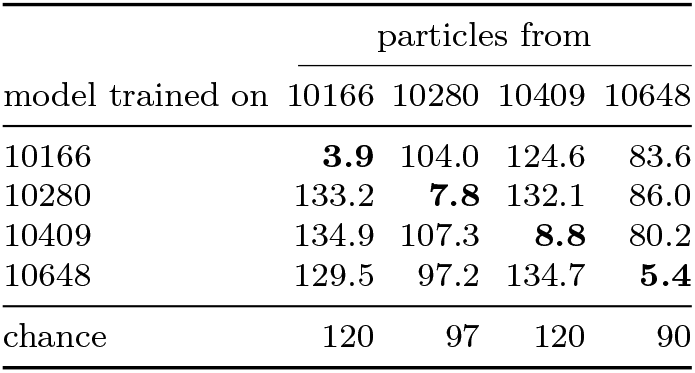
cryoPARES transfer. Each row is a model trained on one target; each column the particles it is applied to. Values are median angular error in degrees on the same 8,000 particles per target, symmetry-aware, using the network stage alone — the local-refinement stage projects the training target’s own reference volume and cannot be run across specimens. The diagonal (bold) is the same-specimen setting every other cryoPARES number in this work reports. Chance is 120° for the C1 targets 10166 and 10409, 97° for C2 10280 and 90° for D2 10648, so every off-diagonal entry is at chance.

## Appendix E Reproducibility settings

**TABLE S17.**
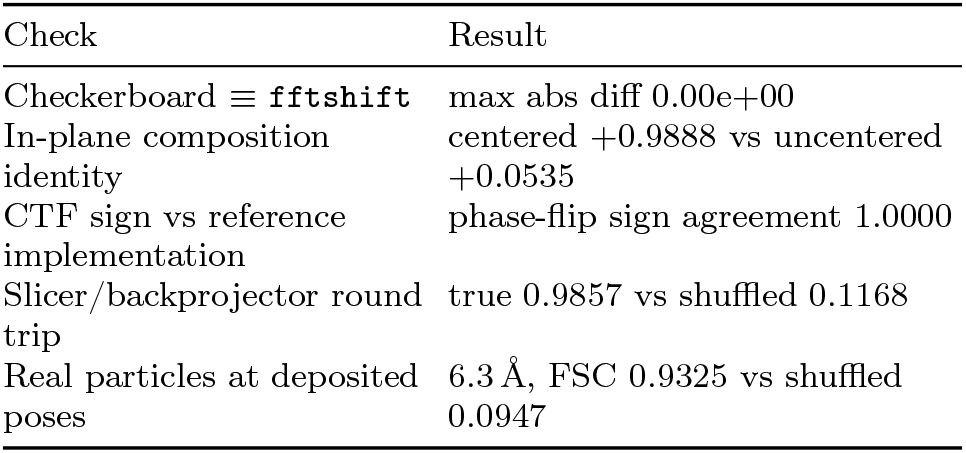
Geometric and CTF conventions checked against external references; all five agree. Real particles back-projected at their deposited poses reconstruct to the *L* = 64 Nyquist limit with mean FSC 0.9325 over shells 3–20 against a shuffled-pose floor of 0.0947; the external anchor is an independent codebase’s reconstructor at 4.17 Å from the same poses and particles.

**TABLE S18.**
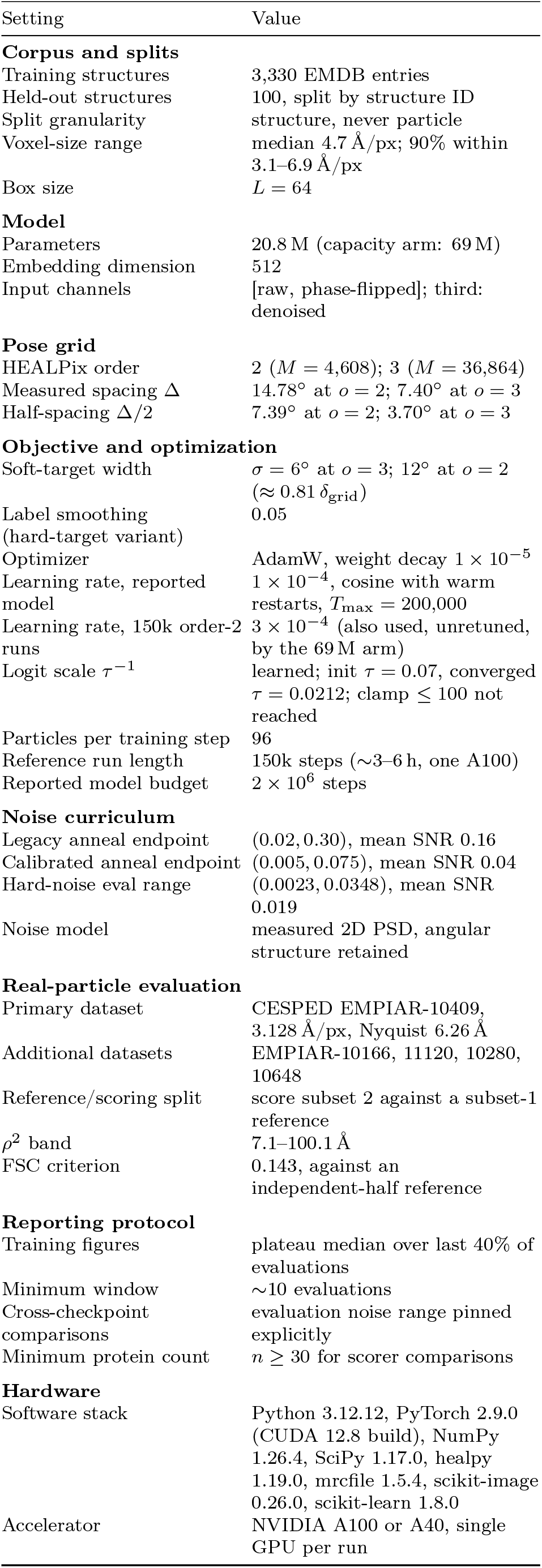
Settings for the reported model and its evaluation.

