## Supplementary Data for "ARCHER: Amortized cross-specimen pose estimation for cryo-electron microscopy"

**TABLE S1.** Notation. Each symbol carries one meaning throughout; $L$ is reserved for the box side, $C$ for a channel count, and $\mathcal{H}$ for the contrast transfer function, following the convention of Refs.[8,67].

| Symbol | Space | Meaning |
|---|---|---|
| *Forward model* (Sec. III A) | | |
| $L$, $C$ | $\mathbb{N}$ | box side in voxels; encoder input channels ($C = 2$) |
| $V$, $\hat{V}$ | $\mathbb{R}^{L\times L\times L}$, $\mathbb{C}^{L\times L\times L}$ | volume and its Fourier transform |
| $y$, $\eta$ | $\mathbb{R}^{L\times L}$ | particle image; its additive noise |
| $\mathbf{R}$ | SO(3) | particle orientation |
| $\xi$, $\mathcal{H}_\xi$ | $\mathbb{R}^3$, $\mathbb{R}^{L\times L}$ | per-particle defocus pair and astigmatism angle; the transfer function they define |
| $\mathcal{S}_\mathbf{R}$ | $\mathbb{C}^{L\times L\times L} \to \mathbb{C}^{L\times L}$ | central-slice operator |
| $\mathbf{q}$, $\mathbf{p}$ | $\mathbb{R}^3$ | Fourier coordinate in slice plane $\Pi_\mathbf{R}$; $\mathbf{p} = \mathbf{R}^{-1}\mathbf{q}$ |
| $k$, $\sigma^2(k)$ | $\mathbb{R}_{\geq 0}$ | shell radius $\|\mathbf{q}\|$; noise power in that shell |
| *Information geometry* (Sec. III B) | | |
| $\hat{e}$, $\hat{e}_j$ | $\mathbb{S}^2$ | rotation axis; the three body axes, $j = 1, 2, 3$ |
| $\delta$, $\boldsymbol{\delta}$ | $\mathbb{R}$, $\mathbb{R}^3$ | rotation perturbation, as angle and as body-axis vector |
| $\mathcal{I}$, $\mathcal{I}_k$ | $\mathbb{R}_{>0}$ | Fisher information of a rotation component; its shell-$k$ part |
| $P_k$, $n_k$ | $\mathbb{R}_{>0}$, $\mathbb{N}$ | signal power in shell $k$; Fourier samples it holds |
| $R_g$, $\kappa$ | $\mathbb{R}_{>0}$ | radius of gyration about $\hat{e}$; geometric factor in Eq. (8) |
| $\Lambda_k$, $c_k$ | $\mathbb{R}$ | shell-resolved score curvature; its correlation at the true pose |
| $\psi$, $\rho^2$ | $[0, \pi]$, $\mathbb{R}_{\geq 0}$ | angle between $\hat{e}$ and $\mathbf{p}$; matched-filter statistic |
| *Estimator* (Secs. III C, III E) | | |
| $\mathcal{G}$, $\Delta$ | $\subset$ SO(3) | orientation grid, $\lvert\mathcal{G}\rvert = 36{,}864$; its mean neighbor spacing |
| $M$, $B$, $N_\text{step}$ | $\mathbb{N}$ | training structures (3,330); particles per training step (96); training steps ($2 \times 10^6$) |
| $\mathbf{t}_i$ | $\mathbb{R}^{L\times L}$ | template rendered at $\mathbf{R}_i \in \mathcal{G}$ |
| $f_\theta$, $f_\phi$ | $\to \mathbb{S}^{511}$ | particle and template encoders (unit-norm, 512-dim) |
| $u_b$, $v_i$ | $\mathbb{S}^{511}$ | particle and template embeddings |
| $s(\mathbf{R})$, $\tau$ | $\mathbb{R}$, $\mathbb{R}_{>0}$ | pose score, Eq. (11); learned temperature |
| $\zeta$ | $(0, 1]$ | orientation information retained by phase flipping, Eq. (12) |
| $\sigma_q$, $\epsilon$ | degrees | geodesic soft-target width; Newton probe angle |
| $g_j$, $\Lambda_j$ | $\mathbb{R}$ | first and second derivatives of $s$ along $\hat{e}_j$ |
| $\mathbf{g}$, $\boldsymbol{\Lambda}$ | $\mathbb{R}^3$, $\mathbb{R}^{3\times 3}$ | gradient and Hessian of $s$ in the chart at $\mathbf{R}$ |
| $m$ | $\mathbb{N}$ | refinement iteration |
| *Denoising channel* (Sec. III D) | | |
| $t$, $T$ | $\{0, \ldots, T-1\}$ | noise-level index; number of levels |
| $\hat{y}_0$, $y_t$, $\varepsilon_t$ | $\mathbb{C}^{L\times L}$ | clean, noised image; its noise ($\varepsilon_{T-1} = \eta$) |
| $\mathbf{A}_t$, $w_t$, $\bar{\alpha}_t$ | $\mathbb{R}^{L\times L}$, $[0, 1]$ | transfer operator $\mathbf{1} \to \mathcal{H}$; its weight; noise schedule |
| $\varepsilon_\vartheta$, $\vartheta$ | $\to \mathbb{C}^{L\times L}$ | denoising U-Net and its parameters (distinct from $\theta$, $\phi$) |
| $\lambda$, $\gamma^\star$ | $\mathbb{R}_{>0}$ | Wiener regularizer $\sigma_n^2/\sigma_s^2$; the gain it induces |
| $\mathcal{L}_\text{CE}$, $\mathcal{L}_\varepsilon$ | $\mathbb{R}$ | pose cross-entropy; $\varepsilon$-prediction loss, Eq. (14) |

### Appendix A: Benchmark scoring details

*Matched particles* (Fig. 5A). Each estimator is scored on data it did not train on. Our encoder and the classical filter never see any of these proteins, so all 8,000 particles per target are held out for both; cryoPARES, being trained per specimen, is scored on the subset lying outside its own training split. Restricting our two arms to that same subset moves their medians by at most 0.09°, so the curves stay comparable.

*Error bars* (Fig. 5B). Bootstrap over particles, 200 resamples.

*Shell correlation* (Fig. 5C). Every arm is reconstructed from the same number of particles per half-set. The band is one standard deviation across targets. The method curves are measured against the deposited map, whereas *half vs half* is our own two half-sets against each other.

*Latent spread* (Fig. 6A). Each grey bar spans the strongest to the weakest canonical direction at that dimension and the dots are the individual directions, ranked. The spread is structure, not uncertainty — RECOVAR is deterministic given its inputs — and it shows that the decline in the mean is the trailing directions decorrelating rather than the leading one degrading. The dashed line is the shuffled-pose floor.

*View-sphere coverage* (Fig. 6B). The 20,000 EMPIAR-10076 particles populate 637 of the 648 cells.

*Ordering by mean error* (Fig. 6C). cryoPARES carries the largest mean error of any arm shown while still producing the best map, because its errors are tail-like rather than diffuse.

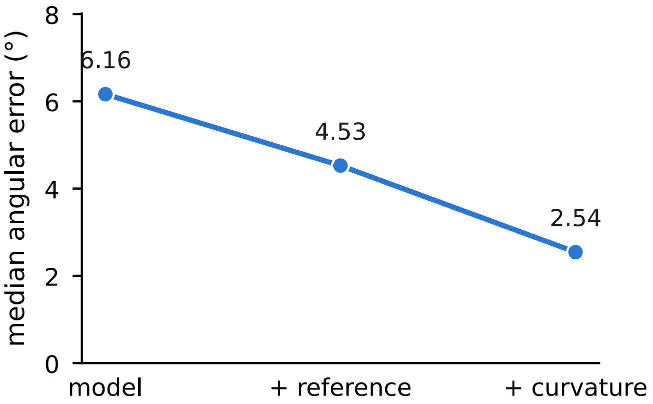


FIG. S1. **Where the accuracy is gained.** Median angular error on EMPIAR-10076 through the refinement ladder: the grid maximum of the learned posterior, re-scoring against a reference rebuilt from those poses, and the curvature step.

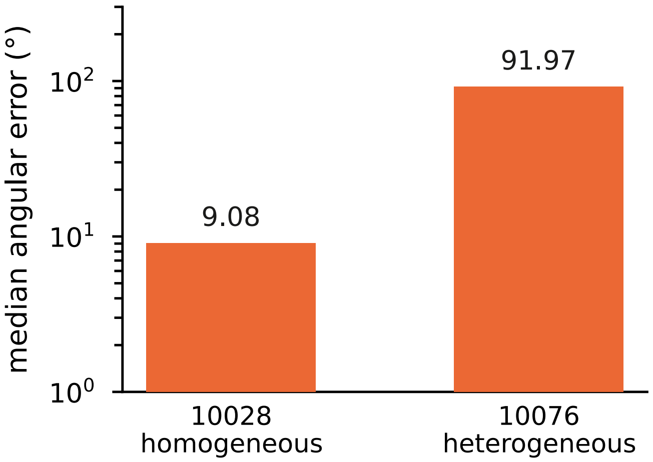


FIG. S2. **Set-size control for the ab-initio comparison.** CryoFastAR evaluated on the homogeneous EMPIAR-10028 at the same particle count used for EMPIAR-10076, isolating specimen heterogeneity from set size.

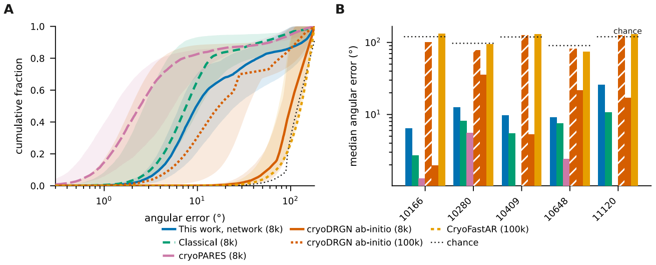


FIG. S3. **Reference-free and per-specimen baselines on matched particles.** (A) Cumulative angular error, median over the five CESPED targets with the full range shaded. (B) Median per target. All arms score the same 8,000 particles per target from one seeded selection; cryoPARES is matched to them by particle identity (`rlnImageName`). Chance is drawn per target, since a point group of order $|G|$ lets a random pose fall nearer an equivalent copy: 120° on the three $C_1$ targets, 97° on the $C_2$ target, 90° on the $D_2$ target. cryoPARES was trained for four of the five targets and reaches 1.29, 5.59 and 2.44° on the matched particles. EMPIAR-11120 carries no cryoPARES bar because no model exists for it; EMPIAR-10409 carries none in this panel alone, where the name-matched subset was not computed, and its model reaches 5.42° on the unmatched 8,000 (Tables S14, S15, S16). The two reference-free methods solve a harder problem — no reference and no translations — and ran at their default settings, each additionally given a solved global rotation and a choice of handedness beyond the point-group quotient every arm receives. Those allowances can only help them, and a random-pose control through the identical alignment stays at 119.6°, so the alignment manufactures no accuracy. Particle count decides cryoDRGN. At 8,000 particles it sits at chance on every target; at 100,000 it reaches 1.96, 5.32 and 17.09° on the three $C_1$ targets, ahead of our network on all three, breaking from chance only near epoch 14 of 30. It reaches 35.49 ($C_2$) and 21.53° ($D_2$) on the symmetric targets, well above chance but far behind the other arms, as `abinit_homo` exposes no symmetry option. None of these transfers: each is a separate optimization costing 2.6–5.3 GPU-hours. CryoFastAR does not improve with count over the same range.

### Appendix B: Architecture and objective

#### 1. Encoders

Both encoders in Eq. (11) are residual convolutional networks[73] with group normalization[92] and SiLU activations, mapping $(B, C, L, L) \to (B, 512)$ with an $\ell_2$-normalized output. They share no weights, and share a topology apart from the stem's input channels. The forward pass is, in order:

1. **Channel assembly.** The image branch of the reported model takes $C = 2$ channels, $[y,\ \mathcal{F}^{-1}[\mathrm{sign}(\mathcal{H})\mathcal{F}y]]$; the template branch takes $C = 1$. The denoiser of Sec. III D supplies a third channel. Each channel is divided by its own standard deviation, so per-protein contrast does not enter the similarity. Wiener-corrected and whitened channels are implemented but are not part of the reported configuration.
2. **Stem.** One 5×5 convolution at stride 2, then SiLU. The stride is not cosmetic: one training step encodes hundreds of templates in one backward pass, and a stride-1 stem would hold every one of them at full $64 \times 64$ resolution simultaneously, which at the widths used here reaches gigabyte scale for a single activation tensor. Halving the resolution cuts that fourfold and costs little at $L = 64$, where the particle already spans most of the box.
3. **Residual trunk.** Four stride-2 blocks. Each block is `conv3×3` → `GroupNorm` → `SiLU` → `conv3×3` → `GroupNorm`, added to a shortcut that is the identity when the shape is preserved and a $1 \times 1$ convolution otherwise, followed by SiLU. Group count

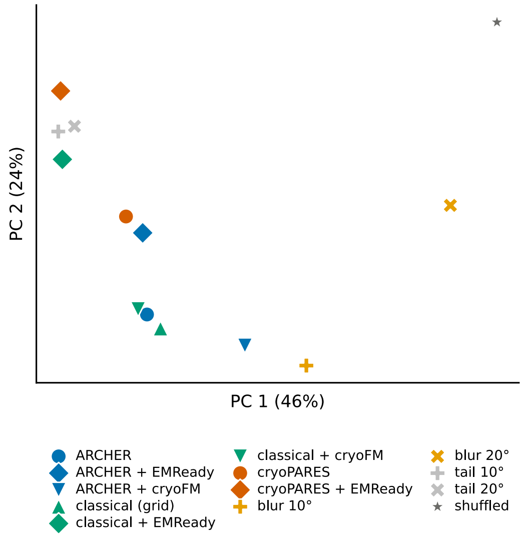


FIG. S4. **Reconstructions of EMPIAR-10409, by method.** Each arm's map of that one specimen, embedded in the same principal-component space as the projections of Fig. 3B; one point is one map. Restored maps group by restorer rather than by the estimator that produced the input, which is the conclusion the displacement cosines of Fig. 3C reach numerically.

is $\min(8, C_{\text{out}})$. Channel width doubles per stage from a base width and is capped at four times that base, so the deepest stages do not dominate the parameter count. For the reported model the base width is 128, giving stage widths $[128, 256, 512, 512]$ and a $2 \times 2$ trunk output at $L = 64$; the five-stage capacity arm of Table S5 reaches $1 \times 1$.

4. **Projection head.** Global average pooling to a vector, then `Linear` $\to$ `SiLU` $\to$ `Linear` to dimension 512, then $\ell_2$ normalization onto $\mathbb{S}^{511}$.

The logit scale $\tau^{-1}$ is a single learned scalar, stored as $\log \tau^{-1}$. Because both embeddings are unit norm, $\langle u, v_k \rangle \in [-1, 1]$, so $\tau^{-1}$ is the only quantity setting how peaked the posterior in Eq. (11) can become. It is initialized at $\tau = 0.07$ and converges to $\tau = 0.0212$ ($\tau^{-1} = 47.1$), a factor of 3.3 sharper than initialization — so the scale is genuinely fitted rather than inherited. A clamp at $\tau^{-1} \leq 100$ guards against an early saturating softmax stalling the gradient; it is never reached.

TABLE S2. The denoised channel, step-matched. Both arms share every setting and differ only in the added third channel. Means over the last 150 step-matched held-out evaluations, spanning steps 600,000–1,345,000; the lead column counts evaluations at which the 3-channel arm is ahead. Consecutive evaluations of one run are correlated, so those counts describe the trajectory and are not independence tests. The accuracy figures reported in the main text are the 2-channel model.

| | 3-channel | 2-channel | leads |
|---|---|---|---|
| Median error | **5.07°** | 5.22° | 122/150 |
| Within 5° | **0.493** | 0.474 | 121/150 |
| Within 15° | 0.755 | 0.753 | 85/150 |

### 3. Objective and optimization

Training minimizes the soft cross-entropy of Eq. (11). The soft-target width is tied to the grid: $\sigma_q = 6°$ for the reported order-3 model on its 7.40° spacing, and $\sigma_q = 12°$ on the 14.78° order-2 grid, i.e. $\sigma_q \approx 0.81\,\Delta$ in both cases. Where a hard target is used instead, label smoothing of

0.05 is applied, matching the reference SO(3)-classifier implementation.[28,93]

Optimization uses AdamW with decoupled weight decay[94] of $1\times10^{-5}$, held constant across every checkpoint in this work, on a cosine schedule with warm restarts.[95] The reported model trains at a learning rate of $1 \times 10^{-4}$ with $T_{\max} = 200{,}000$. The short 150k-step order-2 runs of Table S5 use $3\times10^{-4}$; that value was inherited by the 69 M capacity arm from its 20.8 M control rather than retuned for it, which is the stated scope limit on that comparison.

*a. One schedule property worth stating.* Setting `T_max` equal to the step budget anneals the learning rate to zero exactly while the model is still improving, so a fixed-budget figure is a lower bound set by that budget rather than a converged value. Reported figures are correspondingly plateau medians over a stated window, and any continuation restarts the schedule.

### Appendix C: Full ablation and sweep tables

Throughout this section **bold** marks the best value in a column wherever the column holds directly comparable entries; tables that locate an operating point rather than compare methods carry no emphasis, and the setting adopted is named in the caption.

#### 1. Which configuration produced which number

Table S3 states, for every accuracy figure reported anywhere in this work, the grid order, the evaluation noise range, the particle count, the number of proteins and whether point-group symmetry was quotiented. One difference between configurations dominates the rest. The main-text held-out figures are measured on the order-3 grid (36,864 cells, $\Delta = 7.40°$, soft target $\sigma_q = 6°$), while the ablation and curriculum tables below (Tables S4–S6 and S12) are sweeps over training choices and were run on the cheaper order-2 grid (4,608 cells, $\Delta = 14.78°$, $\sigma_q = 12°$). Halving the grid spacing roughly halves the median error, so the order-2 medians of 10–12° and the order-3 median of 5.0° are one model family measured at two resolutions. Nothing in this work compares an order-2 number against an order-3 one.

#### 2. The anneal-endpoint dose–response

Table S4 gives the per-stage detail behind Sec. III A. The hard-noise metric improves monotonically across all five stages, and the legacy metric across the first four before flattening (0.6141 to 0.6130 at the last), so the trend is a dose–response rather than a single point.

TABLE S3. Every reported accuracy figure and the configuration that produced it. *Grid* is the HEALPix order and the resulting cell count; *SNR* is the evaluation noise range, pinned explicitly whenever checkpoints are compared; *sym.* is whether the geodesic is minimized over the point group. The order-2 rows are training-choice sweeps, not the reported model.

| Figure | Where | Grid | SNR | part. | prot. | sym. |
|---|---|---|---|---|---|---|
| *Held-out synthetic proteins* | | | | | | |
| median 5.0°; < 15° 0.77; top-1 0.41 | §IV A | 3 (36,864) | 0.02–0.30 | 768 | 100 | no |
| < 15° 0.6807; median 10.07° | Table S4, S5 | 2 (4,608) | 0.02–0.30 | 960 | 100 | no |
| hard-noise < 15° 0.4047 | Table S4 | 2 (4,608) | hard | 960 | 100 | no |
| < 15° 0.5463; sym-aware 0.6180 | Table S12 | 2 (4,608) | 0.02–0.30 | 192 | 100 | both |
| classical < 15° 0.6412–0.9161 | Table S9 | 2 (4,608) | swept | 192 | 30 | no |
| *Experimental particles* | | | | | | |
| median 2.5° (EMPIAR-10076) | §IV A | 3 (36,864) | — | 20,000 | 1 | $C_1$ |
| CESPED transfer | Fig. 5A | 3 (36,864) | — | 8,000 | 5 | yes |
| $\rho^2$, < 15° 0.8217 | Table S7 | 2 (4,608) | — | 2,000 | 1 | no |

TABLE S4. Re-annealing to the calibrated endpoint. The legacy metric is held at the pinned $(0.02, 0.30)$ range in every row, so the rows are directly comparable. Stages are numbered along the geometric anneal (ratio $\approx 0.665$ per stage); the warm start enters at stage 6. The stage-10 row is a plateau median over the 12 evaluations of that stage (steps 350k–377.5k). The cycle-2 row continues the stage-10 arm through a learning-rate restart rather than standing beside it, so it is set below the rule; **that row is the reported model**.

| Stage | mean SNR | legacy < 15° | median | hard-noise < 15° |
|---|---|---|---|---|
| Warm start (167.5k) | 1.60 | 0.4688 | 16.8° | — |
| 6 | 0.206 | 0.5042 | 14.7° | 0.1719 |
| 7 | 0.137 | 0.5573 | 12.9° | 0.2255 |
| 8 | 0.091 | 0.5979 | 11.7° | 0.2766 |
| 9 | 0.060 | 0.6141 | 11.4° | 0.3135 |
| 10 (target, plateau) | 0.040 | 0.6130 | 11.4° | 0.3260 |
| *After the cycle-2 learning-rate restart* | | | | |
| Cycle 2 plateau | 0.040 | **0.6807** | **10.07°** | **0.4047** |

#### 3. All three arms at their own plateaus

Table S5 reports the three endpoint arms, each at its own plateau criterion. Table S6 is the controlled version of the same question: there both arms resume the *same* checkpoint under the same schedule and differ only in the SNR bounds.

TABLE S5. Endpoint arms at their own plateau criteria. Arm B is a from-scratch replication at the calibrated endpoint; Arm C is warm-started at the same endpoint; Arm A is warm-started at the legacy endpoint. Bold marks the best of the three. The reported model continues Arm C through a learning-rate restart and exceeds all of them (Table S4, last row).

| | B (scratch) calibrated | C (warm) calibrated | A (warm) legacy |
|---|---|---|---|
| Legacy < 15° | **0.6542** | 0.6130 | 0.5922 |
| Legacy < 30° | **0.7219** | 0.6849 | 0.6781 |
| Median error | **10.5°** | 11.4° | 12.0° |
| Hard-noise < 15° | **0.3760** | 0.3260 | 0.2651 |
| Total steps | 462.5k | 377.5k | 397.5k |
| Real-particle < 15° | **0.4358** | 0.3938 | 0.2727 |

TABLE S6. The controlled endpoint comparison. Both arms resume the *same* checkpoint with the same stage count, caps, plateau window, patience, learning-rate schedule and plateau metric; the only difference is the SNR bounds. Medians over the last 10 matched-step evaluations, from 32 matched steps spanning 170k–247.5k. The calibrated arm leads at 32 of 32 matched steps on both metrics. Consecutive evaluations of one run are correlated, so that count is reported as a description of the trajectory and not as an independence test.

| Endpoint | legacy $< 15°$ | hard-noise $< 15°$ |
|---|---|---|
| Control, legacy $(0.02, 0.30)$ | 0.5156 | 0.1677 |
| Calibrated $(0.005, 0.075)$ | **0.5573** | **0.2255** |
| Difference | +0.0417 | +0.0578 |

#### 4. Detectability of real versus synthetic particles

Table S7 is the measurement establishing that experimental particles carry ample pose information at this box size. A single real particle sits 1.7× above the $2 \ln M = 16.9$ detection threshold and the median particle's true pose ranks first of 4,608.

TABLE S7. Whitened matched-filter detectability, CESPED EMPIAR-10409 at $L = 64$ (3.128 Å/px, Nyquist 6.26 Å) against a reconstructed reference. *rank* is the rank of the true pose among $M = 4{,}608$ candidates for the median particle. The real-particle row is the anchor the noise calibration targets; the synthetic rows bracket it. Real particles are `rlnRandomSubset == 2` scored against the half-1 map, so none contributed to the reference; the partly-overlapping variant gives 0.8380 at $\rho^2 = 29.44$, a 2% difference.

| Arm | $\rho^2$ | true-pose $z$ | rank | $< 15°$ | median |
|---|---|---|---|---|---|
| Real (independent half) | 28.84 | +4.233 | 0 | 0.8217 | 8.1° |
| Real (shuffled poses) | 3.22 | +0.068 | 2159 | 0.0063 | 129.0° |
| Synthetic, SNR 0.03 | 40.85 | +4.955 | 0 | 0.9527 | 7.8° |
| Synthetic, SNR 0.01 | 13.76 | +3.341 | 2 | 0.5303 | 13.2° |
| Synthetic, SNR 0.003 | 4.14 | +1.962 | 115 | 0.0850 | 120.9° |

#### 5. Rejecting junk particles with the pose confidence

The posterior the pipeline already computes carries a usable quality signal at no extra cost. We score the normalized negative entropy $1+\sum_i p_i \log p_i / \log|\mathcal{G}|$, the same confidence the refinement stage uses, and ask how well it separates real particles from junk. CESPED ships curated particles, so junk is constructed, and the construction is the substance of the test: *phase scramble* randomizes Fourier phases at the measured amplitude spectrum, so every shell carries identical power and only structure distinguishes it; *pure noise* redraws amplitudes from the mean shell profile, giving an image with no particle in it; *off-center* translates a real particle 25–45% of the box, the bad pick that actually populates real datasets. Separation is reported as AUROC, the probability that a real particle outscores a junk one. The reference is built from half 1 and the scored particles are drawn from half 0, so the confidence cannot be reading back its own map.

TABLE S8. Junk rejection on EMPIAR-10409, 2,000 particles at $L = 64$ against a half-1 reference, for the reported model. AUROC of the pose confidence, real versus each junk class; 0.5 is no separation. Median confidence is 0.364 on real particles against 0.16–0.18 on junk. The phase-scrambled row is the strongest control, since it matches the real amplitude spectrum shell for shell.

| Junk class | AUROC |
|---|---|
| Phase scramble (matched spectrum) | 0.9626 |
| Pure noise | 0.9690 |
| Off-center pick | 0.9630 |

The reported model separates all three classes at AUROC 0.963–0.969. The phase-scrambled control is the informative one: an image with the real power spectrum and destroyed structure is rejected as readily as pure noise, so the score is responding to particle structure rather than to a difference in image statistics that any variance threshold would catch.

#### 6. The behavioral calibration sweep and the symmetry split

Table S9 gives the sweep behind Sec. III A. Averaged over all held-out proteins the classical scorer settles near the ceiling that ambiguity alone implies (Table S10), well short of the 0.8217 it achieves on real particles. Restricted to non-symmetric proteins it reaches 0.8222 at SNR 0.04, matching the real-particle anchor to within 0.0005. The averaged wall is therefore a property of pooling proteins whose poses are not separable, not of the scorer.

TABLE S9. Classical $< 15°$ against synthetic SNR, split by whether the protein admits indistinguishable poses. The all-proteins column saturates at the ceiling implied by ambiguity alone; the non-symmetric column crosses the real-data anchor of 0.8217 at SNR 0.04. Bold marks that row, the calibrated endpoint adopted throughout; it is the operating point, not the largest entry.

| Synthetic SNR | all proteins | non-symmetric | symmetric |
|---|---|---|---|
| 0.30 | 0.7094 | 0.9161 | 0.2490 |
| 0.08 | 0.6905 | 0.8960 | 0.2317 |
| **0.04** | 0.6412 | 0.8222 | 0.2141 |
| 0.02 | 0.5115 | 0.6361 | 0.1700 |

#### 7. Symmetry: the ambiguity audit and its point-group verification

Table S10 gives the ambiguity statistics on the 30-protein sweep set, where the predicted ceiling and the observed classical saturation agree to 0.2%. Table S11 gives the quantization check on the full 100-protein held-out set. Every one of the 26 symmetric proteins lands on an integer point-group order with none scattered in between, which a correlation artefact could not produce; the dominance of C2 and D2/C4 is what an EMDB-derived corpus of oligomeric complexes should look like.

TABLE S10. Pose ambiguity measured from the volumes alone — no particles, no noise, no scorer.

| Statistic (30-protein sweep set) | Value |
|---|---|
| Mean fraction of poses that are ambiguous | 0.3000 |
| Proteins with $> 50\%$ ambiguous poses | 9 of 30 |
| corr(ambiguous fraction, classical accuracy) | $-0.8431$ |
| corr(voxel size, classical accuracy) | $+0.2922$ |
| Ceiling implied by ambiguity alone | 0.7000 |
| Classical accuracy actually observed | 0.6985 |
| *Extended to all 100 held-out proteins* | |
| Symmetric proteins | 26 |
| Proteins at exactly zero ambiguity | 71 |
| Proteins in between | 3 |
| Achievable ceiling | 0.7388 |
| corr(ambiguity, voxel size) | 0.0124 |

TABLE S11. Independent verification of the ambiguity measurement by point-group quantization. A point group of order $|G|$ predicts exactly $|G| - 1$ indistinguishable alternatives per pose.

| Indistinguishable alternatives | Proteins | Implied $\lvert G\rvert$ | Point group |
|---|---|---|---|
| 1 | 10 | 2 | C2 |
| 3 | 12 | 4 | C4 / D2 |
| 6 | 1 | 7 | C7 |
| 7 | 3 | 8 | C8 / D4 |

TABLE S12. Re-scoring both checkpoints with symmetry separated out. All 100 held-out proteins, 192 particles each, both checkpoints scored at the *same* pinned $(0.02, 0.30)$ range. Arrows run from the 167.5k checkpoint to the plateaued calibrated checkpoint. The non-symmetric row is the control confirming the symmetry-aware correction is not simply inflationary: it moves almost nothing there.

| Subset | $n$ | $< 15°$ | symmetry-aware $< 15°$ |
|---|---|---|---|
| All held-out proteins | 100 | 0.4223 → **0.5463** | 0.4771 → **0.6180** |
| Non-symmetric only | 74 | 0.5255 → **0.6779** | 0.5393 → **0.6892** |
| Symmetric only | 26 | 0.1284 → **0.1719** | 0.3001 → **0.4155** |

#### 8. Per-target accuracy against the per-specimen estimator

Table S13 is the per-target breakdown behind the comparison in Sec. IV A. cryoPARES is scored only where it did not train, and all four arms are scored on exactly those particles, so no arm is credited for particles another arm never saw. Pooling the table by taking the median across the four targets gives medians of 9.5° for the network, 7.5° for our pipeline, 6.6° classical and 4.2° for cryoPARES, and fractions within 15° of 0.689, 0.714, 0.809 and 0.805. The two statistics do not induce the same ordering: cryoPARES leads on the median by a wide margin and ties the classical filter on the tail.

TABLE S13. Per-target accuracy on the particles cryoPARES did not train on. All four arms are scored on the *same* particles of each target — the intersection of our seeded 8,000 with the complement of that target's cryoPARES training split, $n$ below — with the same symmetry-aware geodesic, so the columns are directly comparable. Bold marks the best arm per target. cryoPARES holds the best median on three of the four targets while the classical filter holds the largest fraction within 15° on two of them: the two statistics rank the arms differently, which is why both are reported. EMPIAR-10409 is the clearest case: cryoPARES leads on neither statistic there, taking a median within 0.15° of the classical filter while placing 0.179 fewer particles within 15° — a tail close to that of our unrefined grid argmax.

| | EMPIAR-10166 | EMPIAR-10280 | EMPIAR-10409 | EMPIAR-10648 |
|---|---|---|---|---|
| $n$ scored | 1269 | 4562 | 4079 | 1190 |
| *Median angular error (degrees)* | | | | |
| Network (grid argmax) | 6.28 | 12.48 | 9.91 | 9.04 |
| ARCHER pipeline | 2.97 | 10.14 | 7.06 | 7.90 |
| Classical matched filter | 2.63 | 8.13 | **5.54** | 7.60 |
| cryoPARES | **1.46** | **5.79** | 5.69 | **2.61** |
| *Fraction within* 15° | | | | |
| Network (grid argmax) | 0.7896 | 0.5701 | 0.6249 | 0.7538 |
| ARCHER pipeline | 0.8046 | 0.6223 | 0.6506 | 0.7782 |
| Classical matched filter | **0.9409** | 0.7429 | **0.8120** | 0.8067 |
| cryoPARES | 0.8448 | **0.7646** | 0.6330 | **0.8462** |

TABLE S14. Training cost of the per-specimen comparison. cryoPARES[28] is supervised on the target it is applied to, so each entry is a separate optimization; times are single-GPU wall clock for the completed run on each dataset at the scope used here (200,000 particles, 40 epochs). ARCHER is trained once across 3,330 structures and applied to every target without retraining, so its cost does not recur per specimen — for a new target it is an inference pass.

| Target | GPU-hours |
|---|---|
| EMPIAR-10166 | 13.2 |
| EMPIAR-10280 | 22.8 |
| EMPIAR-10409 | 23.3 |
| EMPIAR-10648 | 31.2 |
| Total, four targets | 90.5 |

### Appendix D: Verification of the geometric conventions

Table S17 lists the checks that fix the sign, centering and transfer-function conventions used throughout. Each compares an implementation against an expectation derived outside it — an analytic identity, a reference implementation, or an independent reconstructor — so agreement is evidence rather than self-consistency.

TABLE S15. Reconstruction quality per target against the deposited map, as the unmasked 0.143 crossing. Unmasked is reported because the masked curve does not cross before Nyquist on EMPIAR-10409 and 10648, so a masked number there is a floor rather than a measurement (Fig. 11C). Our poses were predicted for 60,000 particles per half on EMPIAR-10166 and 10280 against cryoPARES's 100,000, so on those two we also rebuilt cryoPARES from the same 60,000 (last column). Matching the count recovers 0.49 and 0.37 Å of the gap and cryoPARES still leads by 1.62 and 1.69 Å, so the deficit there is mostly pose quality and not set size. The reverse control — rebuilding cryoPARES at the *larger* counts we used on EMPIAR-10409 and 10648 — was not run, hence the two dashes; it would widen the gap on those targets rather than narrow it. Entries are rounded to two decimals, which is coarser than the differences discussed in the main text: on EMPIAR-10409 the three values are 3.634 (ours), 3.625 (classical) and 3.477 Å (cryoPARES), a spread of 0.157 Å. The classical arm, reconstructed from the same particles as ours with no learned component, lands within 0.3 Å of our pipeline on every target: the gap to cryoPARES is therefore a property of reference-conditioned refinement at this working resolution and not of the network. On the two targets where we supply *more* particles than cryoPARES the gap is 0.16 Å and 0.22 Å; the former is the agreement quoted in the main text.

| | particles per half | | resolution (Å) | | | |
|---|---|---|---|---|---|---|
| Target | ours | cryoPARES | ours | classical | cryoPARES | cP at our count |
| EMPIAR-10166 | 60,000 | 100,000 | 5.97 | 5.91 | 3.86 | 4.35 |
| EMPIAR-10280 | 60,000 | 100,000 | 6.30 | 6.14 | 4.24 | 4.61 |
| EMPIAR-10409 | 203,000 | 100,000 | 3.63 | 3.62 | 3.48 | — |
| EMPIAR-10648 | 117,478 | 100,000 | 4.27 | 4.56 | 4.05 | — |

TABLE S16. cryoPARES transfer. Each row is a model trained on one target; each column the particles it is applied to. Values are median angular error in degrees on the same 8,000 particles per target, symmetry-aware, using the network stage alone — the local-refinement stage projects the training target's own reference volume and cannot be run across specimens. The diagonal (bold) is the same-specimen setting every other cryoPARES number in this work reports. Chance is 120° for the $C_1$ targets 10166 and 10409, 97° for $C_2$ 10280 and 90° for $D_2$ 10648, so every off-diagonal entry is at chance.

| | particles from | | | |
|---|---|---|---|---|
| model trained on | 10166 | 10280 | 10409 | 10648 |
| 10166 | **3.9** | 104.0 | 124.6 | 83.6 |
| 10280 | 133.2 | **7.8** | 132.1 | 86.0 |
| 10409 | 134.9 | 107.3 | **8.8** | 80.2 |
| 10648 | 129.5 | 97.2 | 134.7 | **5.4** |
| chance | 120 | 97 | 120 | 90 |

### Appendix E: Reproducibility settings

TABLE S17. Geometric and CTF conventions checked against external references; all five agree. Real particles back-projected at their deposited poses reconstruct to the $L = 64$ Nyquist limit with mean FSC 0.9325 over shells 3–20 against a shuffled-pose floor of 0.0947; the external anchor is an independent codebase's reconstructor at 4.17 Å from the same poses and particles.

| Check | Result |
|---|---|
| Checkerboard ≡ `fftshift` | max abs diff 0.00e+00 |
| In-plane composition identity | centered +0.9888 vs uncentered +0.0535 |
| CTF sign vs reference implementation | phase-flip sign agreement 1.0000 |
| Slicer/backprojector round trip | true 0.9857 vs shuffled 0.1168 |
| Real particles at deposited poses | 6.3 Å, FSC 0.9325 vs shuffled 0.0947 |

TABLE S18. Settings for the reported model and its evaluation.

| Setting | Value |
|---|---|
| **Corpus and splits** | |
| Training structures | 3,330 EMDB entries |
| Held-out structures | 100, split by structure ID |
| Split granularity | structure, never particle |
| Voxel-size range | median 4.7 Å/px; 90% within 3.1–6.9 Å/px |
| Box size | $L = 64$ |
| **Model** | |
| Parameters | 20.8 M (capacity arm: 69 M) |
| Embedding dimension | 512 |
| Input channels | [raw, phase-flipped]; third: denoised |
| **Pose grid** | |
| HEALPix order | 2 ($M = 4{,}608$); 3 ($M = 36{,}864$) |
| Measured spacing $\Delta$ | $14.78°$ at $o = 2$; $7.40°$ at $o = 3$ |
| Half-spacing $\Delta/2$ | $7.39°$ at $o = 2$; $3.70°$ at $o = 3$ |
| **Objective and optimization** | |
| Soft-target width | $\sigma = 6°$ at $o = 3$; $12°$ at $o = 2$ ($\approx 0.81\,\delta_{\mathrm{grid}}$) |
| Label smoothing (hard-target variant) | 0.05 |
| Optimizer | AdamW, weight decay $1 \times 10^{-5}$ |
| Learning rate, reported model | $1 \times 10^{-4}$, cosine with warm restarts, $T_{\max} = 200{,}000$ |
| Learning rate, 150k order-2 runs | $3 \times 10^{-4}$ (also used, unretuned, by the 69 M arm) |
| Logit scale $\tau^{-1}$ | learned; init $\tau = 0.07$, converged $\tau = 0.0212$; clamp $\leq 100$ not reached |
| Particles per training step | 96 |
| Reference run length | 150k steps (∼3–6 h, one A100) |
| Reported model budget | $2 \times 10^6$ steps |
| **Noise curriculum** | |
| Legacy anneal endpoint | $(0.02, 0.30)$, mean SNR 0.16 |
| Calibrated anneal endpoint | $(0.005, 0.075)$, mean SNR 0.04 |
| Hard-noise eval range | $(0.0023, 0.0348)$, mean SNR 0.019 |
| Noise model | measured 2D PSD, angular structure retained |
| **Real-particle evaluation** | |
| Primary dataset | CESPED EMPIAR-10409, 3.128 Å/px, Nyquist 6.26 Å |
| Additional datasets | EMPIAR-10166, 11120, 10280, 10648 |
| Reference/scoring split | score subset 2 against a subset-1 reference |
| $\rho^2$ band | 7.1–100.1 Å |
| FSC criterion | 0.143, against an independent-half reference |
| **Reporting protocol** | |
| Training figures | plateau median over last 40% of evaluations |
| Minimum window | ∼10 evaluations |
| Cross-checkpoint comparisons | evaluation noise range pinned explicitly |
| Minimum protein count | $n \geq 30$ for scorer comparisons |
| **Hardware** | |
| Software stack | Python 3.12.12, PyTorch 2.9.0 (CUDA 12.8 build), NumPy 1.26.4, SciPy 1.17.0, healpy 1.19.0, mrcfile 1.5.4, scikit-image 0.26.0, scikit-learn 1.8.0 |
| Accelerator | NVIDIA A100 or A40, single GPU per run |
